# Evolution of regulatory signatures in primate cortical neurons at cell type resolution

**DOI:** 10.1101/2020.07.24.219881

**Authors:** Alexey Kozlenkov, Marit W. Vermunt, Pasha Apontes, Junhao Li, Ke Hao, Chet C. Sherwood, Patrick R. Hof, John J. Ely, Michael Wegner, Eran A. Mukamel, Menno P. Creyghton, Eugene V. Koonin, Stella Dracheva

## Abstract

The human cerebral cortex contains many cell types that likely underwent independent functional changes during evolution. However, cell type-specific regulatory landscapes in the cortex remain largely unexplored. Here we report epigenomic and transcriptomic analyses of the two main cortical neuronal subtypes, glutamatergic projection neurons and GABAergic interneurons, in human, chimpanzee and rhesus macaque. Using genome-wide profiling of the H3K27ac histone modification, we identify neuron-subtype-specific regulatory elements that previously went undetected in bulk brain tissue samples. Human-specific regulatory changes are uncovered in multiple genes, including those associated with language, autism spectrum disorder and drug addiction. We observe preferential evolutionary divergence in neuron-subtype-specific regulatory elements and show that a substantial fraction of pan-neuronal regulatory elements undergo subtype-specific evolutionary changes. This study sheds light on the interplay between regulatory evolution and cell-type-dependent gene expression programs, and provides a resource for further exploration of human brain evolution and function.

**SIGNIFICANCE:** The cerebral cortex of the human brain is a highly complex, heterogeneous tissue that contains many cell types which are exquisitely regulated at the level of gene expression by non-coding regulatory elements, presumably, in a cell-type-dependent manner. However, assessing the regulatory elements in individual cell types is technically challenging, and therefore, most of the previous studies on gene regulation were performed with bulk brain tissue. Here we analyze two major types of neurons isolated from the cerebral cortex of humans, chimpanzees and rhesus macaques, and report complex patterns of cell-type-specific evolution of the regulatory elements in numerous genes. Many genes with evolving regulation are implicated in language abilities as well as psychiatric disorders.

## INTRODUCTION

Among the numerous phenotypic differences between humans and other primates, the most striking are specializations of social and cognitive abilities including language and executive function, such as abstract reasoning, planning, behavioral inhibition, and understanding mental states of others (1). It has been hypothesized that the evolutionary changes associated with the unique features of human cognition reside, primarily, in the neocortex (2). The neocortex contains multiple cell types, including two major classes of neurons, the excitatory glutamatergic (Glu) projection neurons and the inhibitory GABAergic interneurons, which account for about 80% and 20% of all cortical neurons, respectively (3). The specification and maintenance of these neurons are determined by transcriptional programs that are themselves controlled by the activity of gene regulatory elements (GRE), such as promoters and enhancers (4). GREs recruit transcription factors and chromatin modifiers to control the expression of genes in a cell-type-dependent manner (5). Multiple lines of evidence suggest that phenotypic variation among mammals and susceptibility to brain diseases are largely due to changes in GREs rather than protein-coding sequences (6, 7). Indeed, enhancer changes could cause tissue- or cell-type-dependent adaptations without causing pleiotropic effects that are often associated with changes to genes (8, 9). Therefore, to understand the molecular and cellular differences in brain organization between human and other primates better, it is essential to characterize the mechanisms that drive GRE evolution in individual cell types.

Previous studies used chromatin immuno-precipitation followed by sequencing (ChIP-seq) to assess evolutionary changes of GREs marked by covalent histone modifications in bulk brain tissue (10-12). However, information on key evolutionary changes that could affect the human brain thus far remains limited. One reason is that regulatory changes affecting a particular cell type cannot be reliably inferred from data on bulk brain specimens that conflate signals from all cell types. Mixed signals in bulk specimens likely mask signatures of lower abundance cells, e.g., GABA neurons. In contrast to bulk tissue analysis, single-cell ChIP-sequencing techniques that are currently under development lack sufficient coverage to reliably detect regulatory elements (13, 14). To provide insight into the cell-type-dependent changes that underlie evolution of the human neocortex, we used fluorescence-activated nuclei sorting (FANS) to isolate cortical Glu and GABA neuronal nuclei obtained from one of our closest extant relatives, chimpanzee, and a commonly studied nonhuman primate, rhesus macaque, followed by ChIP-seq and RNA-seq analyses of the epigenome and trancriptome (15, 16). We integrated these data with complementary human transcriptomes and ChIP-Seq data from sorted Glu and GABA neurons (17).

We identified numerous GREs that have not been detected in bulk brain specimens, many of which have neuron-subtype-specific and species-specific histone modifications. We found strong evidence of concordant evolutionary changes in expression and epigenetic regulation for ∼200 genes, highlighting the functional importance of regulatory evolution in neuronal subtypes. These include genes involved in opioid signaling and drug abuse (*OPRM1, PENK, SLC17A8)* (18-20) as well as genes associated with language impairments (*ATP2C2, DCDC2)* (21). We also found neuron subtype- and human-enriched regulatory elements in two genes that are considered among the strongest candidates for enabling or facilitating language abilities, *FOXP2* and *CNTNAP2* (22-24), and identified large clusters of human-specific GREs near genes implicated in neuronal function and brain disorders (e.g., *CDH8, ASTN2, CNTNAP4*) (25-27). Furthermore, we demonstrated that cell-type-specific GREs are more likely to change their activity during primate evolution than GREs shared by different cell types, and found that positional conservation of a GRE in different cell types is not always associated with functional conservation. Our findings provide insight into regulatory evolution that is relevant for brain function and disorders, and present a resource for future studies on comparative epigenomics and neuroscience.

## RESULTS

### Epigenomic profiling of Glu and MGE-GABA neurons from primate brains reveals extensive differences in regulatory landscapes between neuronal subtypes

FANS allows separation and acquisition of nuclei from Glu and medial ganglionic eminence-derived GABA neurons (MGE-GABA) from autopsied cortical samples (15, 17) (**Fig. 1A**). MGE-GABA neurons comprise ∼60-70% of all neocortical GABA neurons and have been implicated in schizophrenia, major depression, autism spectrum disorder (ASD), and epilepsy (28, 29). We recently reported RNA-seq transcriptome profiling and ChIP-seq analysis of histone 3 lysine 27 acetylation (H3K27ac) in Glu and MGE-GABA nuclei that were obtained from dorsolateral prefrontal cortex (DLPFC) (17). H3K27ac is a robust marker of active promoters and enhancers (30, 31). DLPFC is the neocortical region that is crucial for cognition and executive function (1). To investigate regulatory changes across primate evolution, we performed H3K27ac ChIP-seq in Glu and MGE-GABA nuclei from 4 male chimpanzee and 4 rhesus macaque DLPFC, and integrated these datasets with our previously obtained data for humans (17) **(Supplementary (S) Table S1)**.

**Fig. 1.**
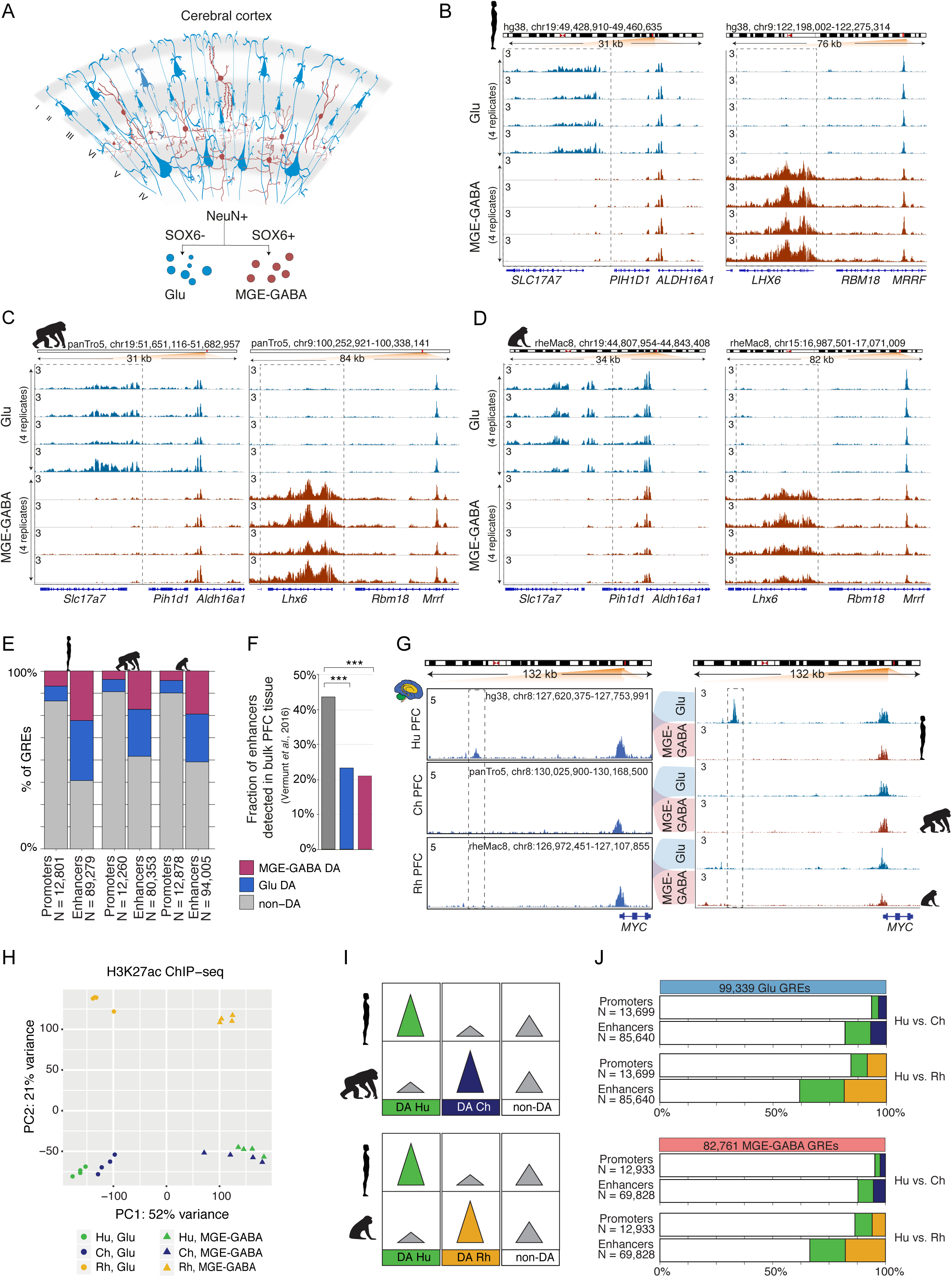
Regulatory changes in Glu and MGE-GABA neurons during primate evolution. A. Schematic of separation of Glu and MGE-GABA nuclei from the DLPFC using FANS. B, C, D. H3K27ac ChIP-seq profiles for human (B), chimpanzee (C), and rhesus macaque (D) in the vicinity of a Glu marker *SLC17A7* (*left panel* in each figure) and an MGE-GABA marker *LHX6* (*right panel* in each figure). Shown are read-per-million (RPM)-normalized ChIP-seq reads for H3K27ac (axis limit 3 RPM). The tracks represent four replicates for each species in Glu (blue) and in MGE-GABA (red) neurons. E. Fractions of promoters or enhancers that are differentially acetylated (DA) or not differentially acetylated (non-DA) between Glu and MGE-GABA neurons in the three species. DA GREs were defined by DESeq2 (FC > 2, FDR < 0.05). F. Fractions of Glu DA, MGE-GABA DA and non-DA human enhancers that overlap GREs in human bulk cortical tissue (10). ***P < 0.0005 by a Fisher’s exact test. G. The regulatory landscapes in the vicinity of the *MYC* locus in the three primate species. Shown are the data in bulk cortex (10) (*left panel*) and in Glu and MGE-GABA neurons (*right panel*). The neuronal analysis confirms the human-specific nature of the enhancer located upstream of *MYC* and further demonstrates the specificity of this enhancer to Glu neurons. Hu, human; Ch, chimpanzee, Rh, rhesus macaque. H. Principal component (PC) analysis of H3K27ac ChIP-seq data for Glu and MGE-GABA neurons from Hu, Ch, and Rh. The non-redundant list of GREs from the 3 species and 2 neuronal subtypes was used for the analysis (**Methods**). Shown are the results for PC1 and PC2. See **Fig. S1F** for the results for PC2 and PC3. I. Schematic of pairwise interspecies comparisons between the ChIP-seq data sets. *Upper panel*: human vs. chimpanzee comparison; *lower panel*: human vs. rhesus macaque comparison. DA, differentially acetylated; non-DA, not differentially acetylated. J. Fractions of promoters or enhancers that are DA or non-DA in pairwise comparisons between species (Hu vs. Ch and Hu vs. Rh). *Upper panel*: Glu GREs. *Lower panel*: MGE-GABA GREs. DA GREs were defined by DESeq2 (FC > 2, FDR < 0.05). The total number of GREs included in the interspecies comparison analysis is shown above each panel.

We produced high-quality H3K27ac ChIP-seq datasets (**Table S2**), with biological replicates showing strong correlation in each cell type and species (all correlation coefficients (r) > 0.8; **Fig. S1A)**. Well-established Glu markers (e.g., *SLC17A7, TBR1*) were enriched in H3K27ac specifically in Glu neurons, whereas typical MGE-GABA markers (e.g., *LHX6, SOX6)* were enriched in MGE-GABA neurons only (**Fig. 1B-D** and **S1B-D)**. We defined putative GREs as regions (peaks) of histone acetylation that were significantly enriched over the background in at least 3 of the 4 donors, detecting a comparable number of GREs (∼100,000) in at least one neuronal subtype in each species (hereafter, Glu or MGE-GABA GREs) (**Methods**; **Table S3**). We then identified GREs that were differentially acetylated (DA) between neuronal subtypes in each species separately, focusing on the subset of GREs with the strongest cell type differences (fold change (FC) > 2, FDR < 0.05) (**Methods, Table S4, Fig. S1E**). We then subdivided GRE regions into proximal H3K27ac peaks (within 1 kb from annotated transcriptional start sites (TSS) in the human genome) which are indicative of active promoters, and distal H3K27ac peaks which are indicative of active enhancers. We confirmed the previously observed greater regulatory diversity of enhancers vs. promoters, detecting significant differential acetylation between cell types in 48-61% enhancers vs. 12-17% promoters (**Fig. 1E**) (9). A substantial proportion of all active neuronal GREs (promoters and enhancres) was differentially acetylated between Glu and MGE-GABA neurons (ranging from 43% to 56% in different species), with a comparable ratio of DA GREs between the neuronal subtypes in each species (**Fig. S1E)**. Gene ontology analysis using GREAT (32) validated the cell type specificity and functional relevance of the regulatory elements we identified (**Table S5**).

To test whether cell-type-specific approach increases the sensitivity and resolution of GRE analysis, we compared the human Glu and MGE-GABA enhancers in the DLPFC with enhancers that were previously identified in bulk human tissues from several brain regions (cortex, cerebellum, and subcortical regions) using H3K27ac ChIP-seq (10, 33). Over 40% of non-DA enhancers but only ∼20% of either Glu or MGE-GABA DA enhancers had been previously identified in bulk cortex (**Fig. 1F**). Despite a higher proportion of Glu vs. MGE-GABA neurons in the cortex, a comparable fraction of Glu DA and MGE-GABA DA enhancers was not detected in bulk cortical tissue. This is likely explained by the large number of unique subpopulations of cortical Glu neurons which were recently uncovered in single nucleus RNA-seq analyses (34). By contrast, the MGE-derived GABA neurons represent a less diverse subset of cortical neurons (34). Our approach also enables the assignment of neuron-subtype specificity for GREs that were previously detected in bulk brain tissues. For example, the human *MYC* enhancer that was previously found to be both human-specific and cortex-specific (10), is exclusively active in Glu but not in MGE-GABA neurons (**Fig. 1G**).

### Substantial regulatory changes in Glu and MGE-GABA cortical neurons during primate evolution

To enable the comparison of the H3K27ac datasets from different species, coordinates of Glu and MGE-GABA H3K27ac-enriched regions from chimpanzee and rhesus macaque were converted to the hg38 human genome assembly using liftOver (10) (**Methods**). Regions with overlapping coordinates were merged (N = 110,270 for Glu; N = 91,560 for MGE-GABA), and liftOver was used again to convert the coordinates to panTro5 and rheMac8. Only GREs that could be mapped onto all three genomes and did not overlap blacklisted regions were included in the subsequent analyses (99,339 Glu and 82,761 MGE-GABA GREs) (**Table S6**; **Methods**). We compared the H3K27ac signals within GREs between the species and neuronal subtypes using Pearson correlation, hierarchical clustering, and principal component analysis (PCA) (**Fig. 1H; Fig. S1F, G; Methods**). Based on correlation and clustering analyses, we detected greater differences between cell types in each species compared with the differences between species in each cell type. Clustering and PCA showed a clear separation of samples into six groups according to cell type and species, with human and chimpanzee clustering closer together vs. rhesus macaque in each cell type.

To investigate the regulatory changes during primate evolution, we analyzed H3K27ac levels within the GREs that were differentially acetylated between species in each neuronal subtype in pairwise comparisons (i.e., species-enriched GREs) (**Methods**; **Fig. 1I, J; Fig. S1H; Table S7**). In both Glu and MGE-GABA, ∼14-16% of promoters (2,146 in Glu and 1,754 in MGE-GABA) and ∼34-38% of enhancers (32,785 in Glu and 23,435 in MGE-GABA) were differentially acetylated in human vs. rhesus macaque (FDR < 0.05, FC >2) (**Fig. 1J**). Consistent with the closer evolutionary relationship between human and chimpanzee, there were fewer DA promoters (896 in Glu and 609 in MGE-GABA) and enhancers (15,625 in Glu and 8,495 in MGE-GABA) between human and chimpanzee. Thus, using a cell-type-specific analysis, we showed that, in each neuronal subtype, ∼30% of the GREs underwent significant changes in primate evolution.

### Neuron-subtype-specific regulatory elements change more frequently during evolution than non-specific ones

Previous work in bulk brain tissues has shown that evolutionary changes preferentially occur in GREs that are mainly active in a single brain region (cortex, midbrain or cerebellum), but not in multiple structures (10). This result fits a model in which changes in GREs that are not region-specific are more likely to cause pleiotropic effects, which might be detrimental to the fitness of the organism and are, therefore, selected against during evolution. We hypothesized that this trend also extends to evolutionary changes in the activity of cell-type-enriched GREs. Only a limited number of GREs have changed between human and chimpanzee, precluding a reliable statistical analysis. We therefore assembled two sets of GREs that were detected in human and/or rhesus macaque in each neuronal subtype (93,222 for Glu and 79,181 for MGE-GABA), and compared cell type specificity of DA and non-DA GREs between the two species (**Fig. 2A**). We found that DA GREs were significantly more often cell-type-specific than non-DA ones (Fisher’s exact test, p-values < 2.2e-16; odds ratios: 2.8 for Glu and 2.2 for MGE-GABA). Conversely, human GREs that were non-DA between neuronal subtypes were significantly less frequently DA between the two species compared to GREs which were DA between Glu and MGE-GABA (Fisher’s exact test, p-values < 1e-4; **Fig. S2A**).

**Fig. 2.**
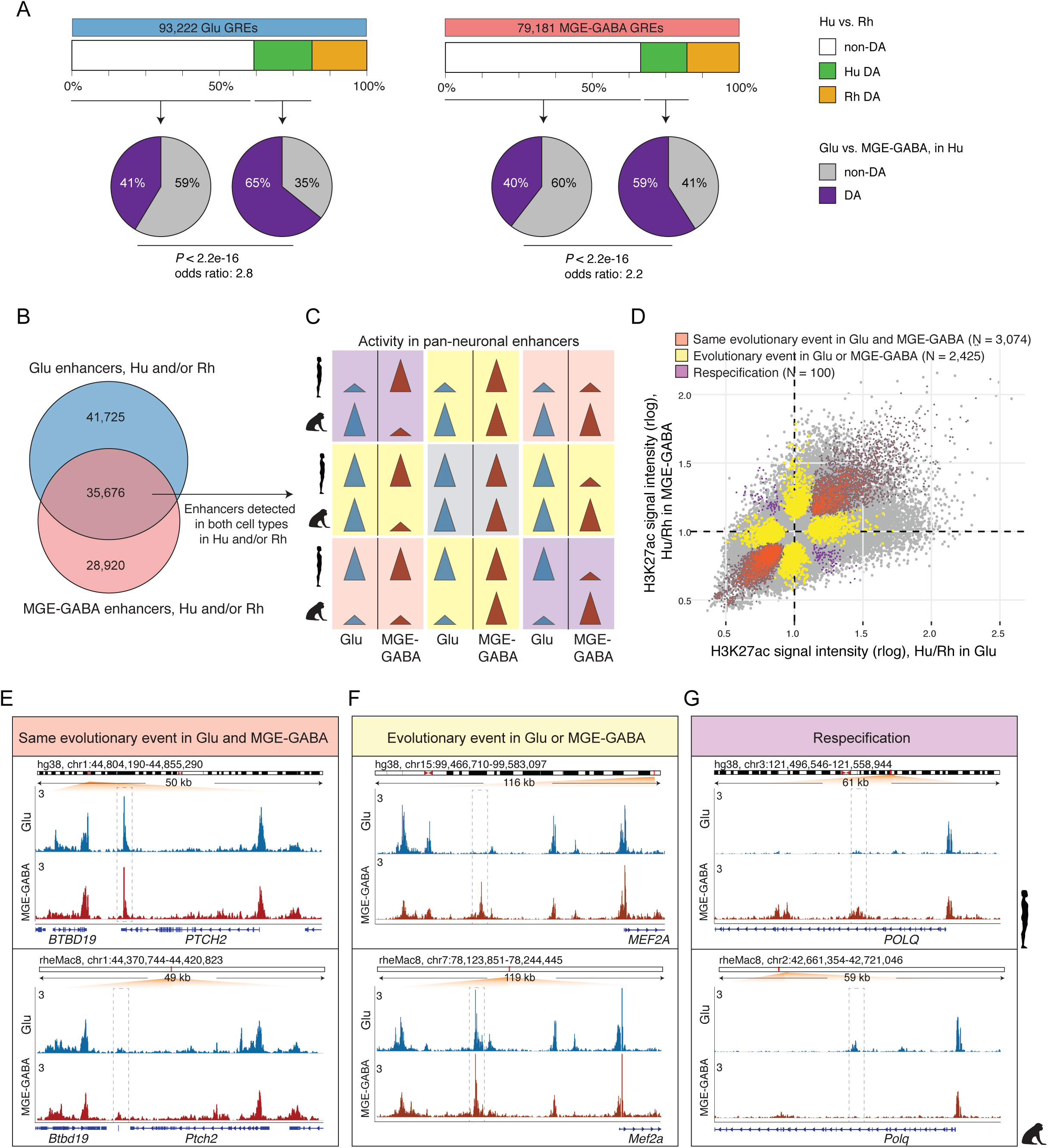
Evolutionary changes in neuron-subtype-specific and pan-neuronal regulatory elements. A. Analysis of cell type specificity of GREs that were DA (Hu>Rh DA, *green boxes*) or non-DA (*white boxes*) between Hu and Rh. *Left panel*: GREs detected in Glu neurons in Hu and/or Rh. *Right panel*: GREs detected in MGE-GABA neurons in Hu and/or Rh. Pie charts indicate the fractions of the GREs that are DA (shown in *grey*) or non-DA (shown in *purple*) between the neuronal subtypes in human. In both neuronal subtypes, Hu>Rh DA GREs were more often neuron-subtype-specific than Hu vs Rh non-DA GREs (p-values <2.2e-16 by Fisher’s exact test). B. Venn diagram showing the overlap between Glu and MGE-GABA enhancers that were identified in Hu and/or Rh. The overlapping enhancers, depicted in *purple*, are positionally shared in Glu and MGE-GABA neurons (pan-neuronal, i.e., active in both neuronal subtypes) in Hu and/or Rh. C. Schematic of possible evolutionary changes in enhancers that are pan-neuronal in Hu and/or Rh. *Gray*: no evolutionary change in any neuronal subtype; *orange*: same direction of evolutionary change in both subtypes; *yellow*: evolutionary change in only one subtype; *purple*: different direction of evolutionary change between subtypes (respecification). D. Simultaneous cross-species and cross-cell-type analysis of H3K27ac signal intensities for pan-neuronal enhancers in Hu and/or Rh. Highlighted are the DA GREs depicted in **Fig. 2C** which were confirmed in the analysis (**Methods, Fig. S2C**) to display *(1)* the same direction of evolutionary change in both neuronal subtypes (*orange*), *(2)* an evolutionary change in only one neuronal subtype (*yellow*) or (3) a respecification change (*purple*). E. Example of an enhancer with the same direction of evolutionary change in Glu and MGE-GABA within the *PTCH2* locus (dashed box). Representative tracks for Glu (blue) and MGE-GABA (red) are displayed for each species. RPM-normalized H3K27ac ChIP-seq reads (axis limit 3 RPM) are presented here and in F-G. F. Example of an enhancer with an evolutionary change in only one neuronal subtype, MGE-GABA, in the vicinity of the *MEF2A* locus (dashed box). G. Example of an enhancer with an evolutionary change of opposite directions in Glu and MGE-GABA neurons (respecification) within the *POLQ* locus (dashed box).

To further validate this finding, we used a threshold-free approach by directly comparing the enrichment of enhancers in neuronal subtypes (e.g. difference between Glu and MGE-GABA in human, *ΔCellType*_*human*_ *= H*3*K*27*ac*_*Glu,human*_ − *H*3*K*27*ac*_*GABA,human*_) vs. the evolutionary change (e.g., difference between human and macaque in Glu cells, *ΔSpecies*_*Glu*_ *= H*3*K*27*ac*_*Glu,human*_ − *H*3*K*27*ac*_*Glu,rhesus*_) (**Methods**). This analysis used independent biological samples to estimate *H*3*K*27*ac*_GLU,*human*_ at each step of the analysis, to avoid spurious correlations driven by noise in individual datasets. Across all enhancers detected in human Glu cells, we found a significant correlation between the cell type specificity and the evolutionary change (Spearman r = 0.203, p < 1e-16) (**Fig. S2B, top row**). No such correlation was observed when the enhancers were randomly shuffled. A similar pattern was observed in human MGE-GABA cells (r = 0.106, p < 1e-16; **Fig S2B, bottom row**). These analyses confirm that evolutionary divergence of neuronal enhancers preferentially occurs in a cell-type-specific manner.

### Pan-neuronal enhancers undergo neuron-subtype-specific evolutionary changes

Whereas GREs active in both Glu and MGE-GABA (hereafter, pan-neuronal GREs) are more often evolutionarily conserved between human and rhesus macaque than cell-type-specific ones, a substantial proportion of pan-neuronal GREs change in at least one neuronal subtype (**Fig. S2A**). Thus, our data offered a unique opportunity to compare the evolution of enhancers that are active in at least 2 closely related cell types. We found that, among these shared enhancers (N = 35,676, dark purple in **Fig. 2B**), 19,225 (∼54%) showed no evolutionary changes (gray panel in **Fig. 2C**), whereas the rest (N=16,451) changed between the species in at least one cell type (DA analyses between human and rhesus macaque in Glu or MGE-GABA, FC > 2; FDR < 0.05; yellow, orange, and purple panels in **Fig. 2C**). We then asked whether evolutionary changes in the latter group were similar in Glu vs. MGE-GABA neurons. We categorized the enhancers into: ***(1)*** enhancers with the same direction of evolutionary change in both Glu and MGE-GABA neurons (orange panels in **Fig. 2C), *(2)*** enhancers with an evolutionary change in only one cell type (yellow panels in **Fig. 2C)**, and **(3)** enhancers with an evolutionary changes of opposite directions in Glu and MGE-GABA neurons. The latter group represents a small subset of enhancers that may be functionally respecified (10, 35), with activity switching from one neuronal subtype in one species to a different neuronal subtype in another species (purple panels in **Fig. 2C**). We used a linear model to further assess the statistical significance of these evolutionary events, assigning events to several categories depending on the proportion of variance explained by the corresponding factor (η^2^) and the FDR value (**Methods, Fig. S2C**). By overlapping the results of this analysis with the DA analysis (**Fig. 2C**), we identified 3,074 enhancers that changed in the same direction in both cell types, 2,425 enhancers that changed in one but not in the other cell type, and 100 enhancers that underwent a respecification (**Fig. 2D, Table S8**).

Examples of these three types of evolutionary events are highlighted in **Fig. 2E-G**. The enhancer in the *PTCH2* locus showed increased activity in human vs. rhesus macaque in both neuronal subtypes (**Fig. 2E**). Another enhancer, located ∼60kb upstream of the *MEF2A* promoter, illustrates an evolutionary change that occurred in only one cell type. The pan-neuronal activity of this enhancer in rhesus macaque is contrasted with strictly Glu-specific H3K27ac signal in human (**Fig. 2F**). Finally, an enhancer in the *POLQ* locus provides a rare example of respecification, as it is Glu-specific in human and MGE-GABA-specific in rhesus macaque (**Fig. 2G**). Altogether, these results indicate that enhancers in conserved genomic positions between species in the same tissue can undergo different evolutionary changes in different cell types. Therefore, positional conservation of an enhancer is not always a reliable proxy for its functional conservation, suggesting that variation of enhancer activity between cell types is most likely greater than can be inferred based on positional conservation alone.

### Concordant evolutionary changes in GREs and gene expression underscore functional relevance of regulatory evolution in neuronal subtypes

We next examined how the evolutionary changes in the regulatory landscape of the neuronal subtypes are reflected in changes in gene expression. We performed RNA-seq on FANS-separated Glu and MGE-GABA nuclei purified from the same DLPFC specimens that were used for the H3K27ac profiling (**Table S1**), obtaining high quality transcriptomes for each species and cell type (**Methods, Figs. S3A-E, Table S9**). In agreement with previous findings in various bulk tissues and species (36), the expression of protein-coding genes (N=16,846) was much more conserved between species than the expression of long intergenic non-coding RNA (lincRNA) genes (N=518) **(Fig. S3F**).

Next, in each neuronal subtype, we identified DA GREs and differentially expressed (DE) genes that were enriched or depleted in a particular species compared to the two other species (**Fig. 3A left panel, Fig. S3G**). We denoted these upregulated or downregulated GREs or genes as species-specific up- or down-DA GREs or DE genes. To focus on high confidence DA GREs and DE genes, we applied conservative FC thresholds in both analyses (FC > 2, FDR < 0.05) (**Methods**; **Tables S10-12**). We found thousands of DA GREs and hundreds of DE genes for each comparison, with the largest numbers of species-specific GREs and genes detected in rhesus macaque (**Fig. 3B, C**). The analysis of DE genes showed that approximately half of the genes that were species-specific in one neuronal subtype were also specific for the same species in another neuronal subtype (**Table S12**). Also, for both neuronal subtypes, the species-specific up-DE genes significantly overlapped with the up-DE genes that have been recently reported for the same species in sorted neuronal (NeuN+) nuclei from the DLPFC (11-31% overlap for Glu DE genes and 12-23% overlap for MGE-GABA DE genes, p-values <2.5e-6 by hypergeometric test; **Table S13**) (37). In addition, our gene expression data were in good agreement with the results of human vs. rhesus macaque DE analysis in bulk adult DLPFC reported by the PsychENCODE consortium (17-19% overlap, p-values < 2.8e-14 by hypergeometric test; **Table S14**) (38).

**Fig. 3.**
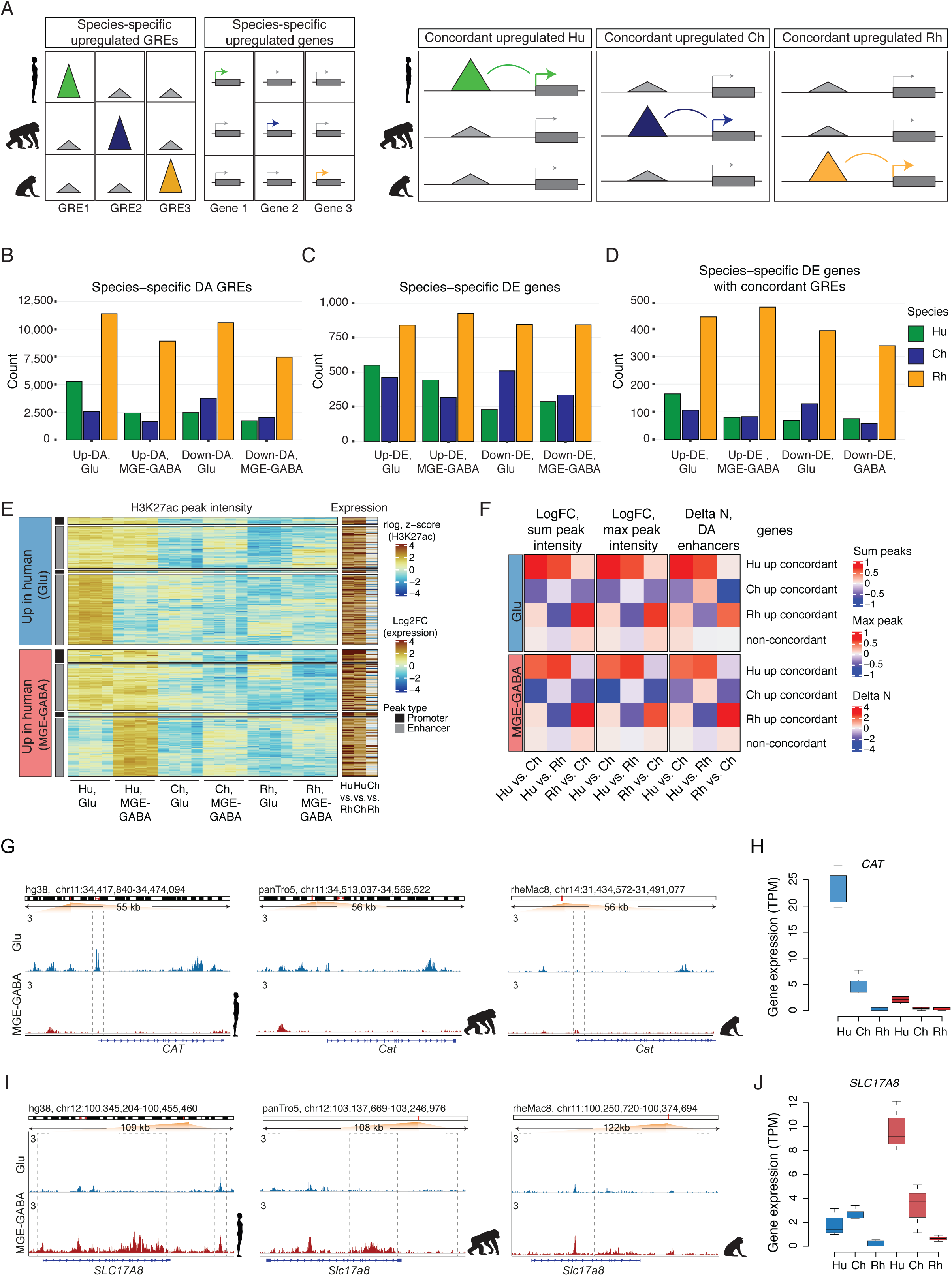
Concordant evolutionary changes in GREs and gene expression. A. Schematic of the analysis of association between species-specific upregulated GREs and genes. *Left panel*: Identification of species-specific DA GREs and species-specific DE genes. *Right panel*: Identification of concordant pairs of species-specific upregulated DA GREs and DE genes. B. Numbers of species-specific upregulated or downregulated DA GREs (up-DA or down-DA GREs) detected for each species in Glu or MGE-GABA neurons. C. Numbers of species-specific upregulated or downregulated DE genes (up-DE or down-DE genes) detected for each species in Glu or MGE-GABA neurons. D. Numbers of species-specific upregulated or downregulated DE genes with concordant species-specific DA GREs. E. Heatmap of H3K27ac signal intensities and interspecies changes in gene expression for concordant pairs of human-specific up-DA GREs and up-DE genes. *Upper panel*: Glu concordant GRE-gene pairs. *Lower panel*: MGE-GABA concordant GRE-gene pairs. K-means clustering of the data sets for H3K27ac signal in individual Hu, Ch, and Rh samples resulted in 2 major clusters for each neuronal subtype, reflecting that ∼50-60% of concordant human up-DA GREs were neuron-subtype-specific, whereas the remaining group was non-DA between Glu and MGE-GABA. Concordant promoter- and enhancer-gene pairs are shown separately for each cluster. Human-specific upregulation of gene expression is significantly stronger for promoter- than enhancer-gene pairs (also see **Fig. S3L**). Heatmap colors indicate normalized H3K27ac signal intensities (rlog) for each replicate sample and log2 fold changes (FC) of normalized RNA-seq reads (Transcripts Per Million, TPM) for interspecies comparisons. Fold changes of gene expression levels between species represent data for Glu neurons (*top panel*) and for MGE-GABA neurons (*bottom panel*). F. Mean interspecies changes of quantitative features of gene regulatory domains. Shown are data for species-specific upregulated concordant genes, as well as for non-concordant genes for: *(1)* the sum of normalized H3K27ac signal intensities (rlog) for all enhancers within a gene regulatory domain, *(2)* the maximum normalized H3K27ac signal intensity among all enhancers within a gene regulatory domain, and (3) the number of species-enriched DA enhancers in pairwise interspecies comparisons. *Upper* and *lower panel*: data for Glu and MGE-GABA neurons, respectively. G. Evolutionary regulatory changes within the *CAT* locus. RPM-normalized H3K27ac ChIP-seq reads are shown here and in **Fig. 3I**. Representative tracks for Glu (*blue*) and MGE-GABA (*red*) neurons are shown here and in **Fig. 3I** for each species. The dashed box highlights a human- and Glu-specific H3K27ac peak in the promoter region of *CAT*. H. Evolutionary gene expression changes of *CAT*. TPM-normalized RNA-seq reads for each cell type (*blue* for Glu neurons and *red* for MGE-GABA neurons) and species are shown here and in **Fig. 3J**. I. Evolutionary regulatory changes within the *SLC17A8* locus (see **Fig. 3G** for details). The dashed boxes highlight GREs which display human-specific upregulation in MGE-GABA neurons. J. Evolutionary gene expression changes of *SLC17A8* (see **Fig. 3H** for details).

We then linked species-specific GREs to nearby genes using GREAT (**Methods**) and overlapped the resulting gene sets with genes showing species-specific expression (**Fig. 3A right panel, Fig. S3H, Tables S15, S16**). This analysis yielded hundreds of genes with evidence of concordant species-specific evolutionary changes in gene expression and in at least one associated GRE (**Fig. 3D**). The fraction of species-specific DE genes which had a concordant DA GRE varied for each species and was in the range of 18-30% in human, 17-26% in chimpanzee, and 40-53% in rhesus macaque for up- or down-regulated DE genes in Glu or MGE-GABA (**Table S17**). We used random shuffling (**Methods**) to assess statistical significance of the observed concordant association between GREs and genes. For all categories of concordant changes (species-specific up- or downregulated, Glu or MGE-GABA neurons, promoters or enhancers), concordant GRE-gene pairs were significantly over-represented compared with random pairs (all p-values < 1e-7) (**Fig. S3I**). Thus, a sizable proportion of the identified concordant GRE-gene pairs likely represents functional associations that underlie evolutionary differences in gene expression between the primate species, rather than resulting from coincidental co-localizations of species-specific DA peaks and DE genes.

Among the concordant human up-DA GREs, ∼50-60% were specific for Glu or MGE-GABA, whereas the remaining group displayed a strong H3K27ac signal in both neuronal subtypes (**Fig 3E**). Separation of GREs into promoters and enhancers showed that the majority of genes with concordant evolutionary changes in expression and promoter signal also had an evolutionary change in at least one enhancer (70% in Glu and 64% in MGE-GABA) (**Fig. S3J**). Lastly, in all 3 species, concordant genes often associated with multiple concordant enhancers (23-45% of human- or chimpanzee-specific and ∼60% of rhesus macaque-specific concordant up- or down-regulated DE genes in Glu or MGE-GABA, **Fig. S3K**).

Whereas changes in promoter signals were associated with larger changes in gene expression compared with changes in enhancers (**Fig. 3E, Fig S3L**), the neuronal regulatory landscapes often encompasse more than one enhancer per gene (4, 39). Therefore, we asked if additional quantitative features that represent aggregate measures of an entire gene regulatory domain (as defined by GREAT with the basal plus extension setting, **Methods**), such as the number of DA regulatory elements per gene (40) or the intensity of an enhancer signal (maximum and sum), could provide independent validation of the functional evolutionary change in a concordant DA enhancer-gene pair. We found that, in both neuronal subtypes, each of these regulatory features significantly correlated with gene expression (**Fig. S3M, N**) and that evolutionary changes in these features were significantly correlated with evolutionary changes in gene expression (Spearman correlation, all p-values < 2.2e-16; **Figs. S3O, P**). Finally, we confirmed that the majority of the concordant enhancer-gene pairs represent genes with an evolutionary change not only in individual enhancer but also in the entire regulatory domain (**Fig. 3F**). These gene sets, with their associated epigenetic features, are compiled in **Table S18**, which provides a resource to enable future hypothesis-driven functional and evolutionary studies on primate brains.

Two representative examples of human-specific concordant GRE-gene pairs are *CAT* and *SLC17A8* loci (**Fig. 3G-J)**. *CAT* encodes the hydrogen peroxide-degrading enzyme catalase. *CAT* is more strongly expressed in Glu vs. MGE-GABA neurons and is preferentially expressed in hominids (i.e., humans and chimpanzees) vs. rhesus macaque, with a significantly higher expression in human vs. chimpanzee. In addition to the strong human-specific H3K27ac enrichment in the promoter, we also detected an enrichment in several Glu-specific *CAT* enhancers in human or chimpanzee vs. rhesus macaque. Accumulation of H_2_O_2_ in oxidative metabolism has been linked to oxidative-stress-associated neurodegenerative disorders (41). Elevated expression of *CAT* in human neurons could, therefore, reflect the evolution of protective mechanisms that limit oxidative damage resulting from the higher metabolic activity of the human brain compared to that of chimpanzee (42-44). *SLC17A8* encodes vesicular glutamate transporter 3 (VGLUT3) and is selectively up-regulated in human MGE-GABA neurons, with corresponding gains in human-specific enhancers. *SLC17A8* is implicated in cocaine abuse (20) as well as in anxiety-related behaviors (45).

### Evolutionary changes in regulation and expression of genes associated with language ability

One of the most striking differences between humans and other primate species is language ability. We examined 10 human genes (*ATP2C2, CMIP, CNTNAP2, DCDC2, DYX1C1, FOXP2, KIAA0*3*19, NFXL1, ROBO1, ROBO2*) that had been associated with language impairment or developmental dyslexia (21, 46). Among them, *ATP2C2*, which is linked to language impairment (47), is strongly expressed in Glu neurons specifically in human (5 or 8 fold change vs. chimpanzee or rhesus macaque, adj. p-values < 1.2e-5), which was concordant with changes in multiple regulatory features (**Table S18**). Also, *DCDC2*, which is implicated in reading proficiency (48), showed concordant upregulation of expression and GRE signals in hominids compared with rhesus macaque, specifically in MGE-GABA neurons.

Whereas the other 8 genes associated with language impairment did not show any concordant evolutionary changes in their regulation and expression, we detected many human-specific (and often cell-type-specific) DA GREs that were linked to several of these genes (**Table S19**). Human-specific enhancers were found in *CNTNAP2* (N = 8 in Glu, N = 6 in MGE-GABA) and *FOXP2* (N=1 in Glu) loci (**Fig. 4A; Fig. S4A-B**). These two genes have been suggested to be required for the proper development of speech and language in humans (22-24). The functional importance of these regulatory changes remains unclear, as we did not observe any significant human-specific changes in gene expression for *FOXP2* or *CNTNAP2*. Nevertheless, it appears plausible that the increase in regulatory complexity engenders context-dependent mechanisms of regulation of *FOXP2* or *CNTNAP2* expression in anatomical (including brain areas and subpopulations of Glu or MGE-GABA neurons), environmental or developmental contexts, which could, in turn, facilitate the emergence of language skills. Notably, cortical areas other than DLPFC, such as inferior frontal cortex and temporal cortex, constitute the core of the brain language network (49, 50). The evolutionary changes in regulation and expression of *FOXP2* or *CNTNAP2* in these cortical areas remain to be investigated.

**Fig. 4.**
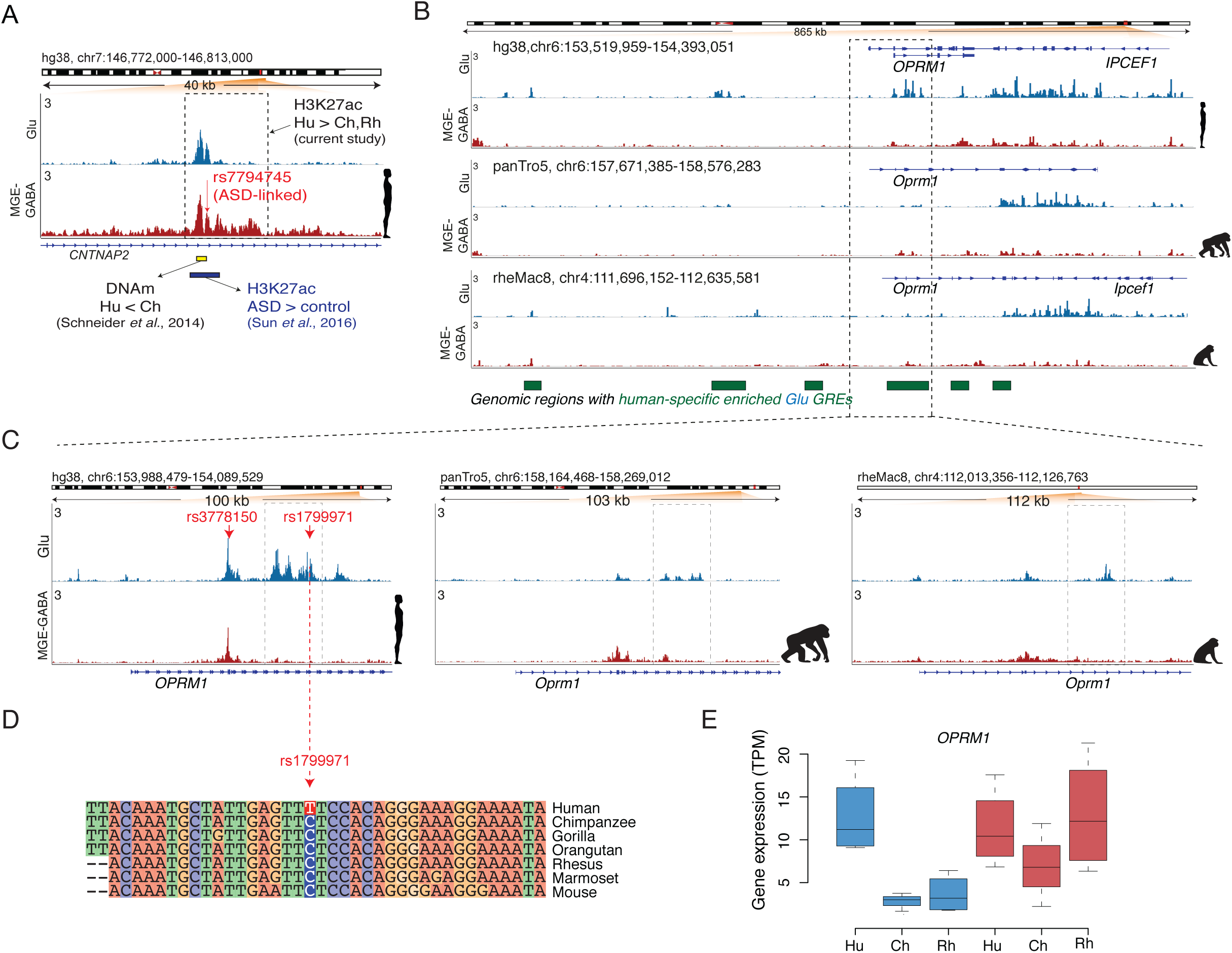
Human-specific upregulated GREs harbor genes implicated in language, ASD and opioid addiction. A. Evolutionary regulatory (H3K27ac and DNA methylation) and ASD-associated changes at an enhancer within the *CNTNAP2* locus in humans (also see **Fig. S4B**). The enhancer (*shown in a dashed box*) is located within the second intron of *CNTNAP2* and shows human-specific upregulation of the H3K27ac signal in MGE-GABA neurons. Shown are: ASD-associated SNP *rs7794745* (*red arrow*) (52), the region with the largest decrease in DNA methylation (DNAm) in Hu vs. Ch (*green bar*) (53), and the position of an H3K27ac peak (*brown bar*) that is upregulated in ASD vs. control subjects (55). B. The *OPRM1* locus shows a high density of Glu human-specific up-DA GREs (the regions marked as *green* boxes depict areas with one or several Hu up-DA GREs). The leftmost region exemplifies an evolutionary respecification change from a Rh-enriched enhancer in MGE-GABA to a Hu-enriched enhancer in Glu. C. H3K27ac profiles within the *OPRM1* locus in 3 species and 2 neuronal subtypes. Evolutionary regulatory changes were found within the 5’ region, including human-upregulated promoter and enhancer GREs. The opioid abuse-associated SNPs *rs3778150* and *rs1799971* (*red arrows*) overlap with a human-specific up-DA promoter and human-specific up-DA enhancer, respectively. D. Genomic alignment of the 40-bp region centered on the *rs1799971* SNP in humans. The human-specific nucleotide substitution at the SNP position (C -> T) is highlighted. Notice a high level of sequence conservation in the immediate vicinity of the SNP. The C nucleotide in humans represents the minor allele, which has been associated with opioid abuse (56). E. Evolutionary changes in gene expression of *OPRM1* in Glu neurons. Gene expression changes were concordant with the regulatory changes, suggesting human-specific and Glu-specific upregulation of *OPRM1*.

A recent DNA methylation study in the PFC of human and chimpanzee identified four regions within the *CNTNAP2* locus that were differentially methylated between the two hominids (51). Remarkably, the region with the largest decrease in DNA methylation in human vs. chimpanzee (region B in that study) overlapped with one of the MGE-GABA- and human-specific upregulated enhancers detected in our study (**Fig. 4A**). Also, an ASD-linked SNP rs7794745 (52), that is located ∼280 bp from differentially methylated region B, colocalizes with the same *CNTNAP2* enhancer (**Fig. 4A**). The major allele nucleotide of rs7794745 (A>T) is a human-specific substitution compared to other primate species (**Fig S4C**). This enhancer was also identified and found to be upregulated in ASD in a recent H3K27ac study in bulk brain tissues, including PFC (**Fig. 4A**) (53). These findings link a cell-type-specific regulatory element that underwent an evolutionary change in humans to a risk locus associated with a neuropsychiatric disorder.

### Human-specific upregulated GREs form neuron-subtype-dependent clusters located near genes associated with neuronal function and neuropsychiatric disorders

We found continguous genomic regions with a high density of human-specific upregulated GREs, which suggested a non-uniform distribution of these GREs across the genome. This trend is exemplified by a ∼1-Mb-long genomic region upstream of *CDH8* on chromosome 16 that contains a cluster of 11 human-specific up-DA GREs in Glu neurons (**Fig. S4D**). Applying stringent statistical analysis (**Methods**), we identified a significant enrichment of human up-DA GREs over the background distribution of all human GREs in 23 clusters, with 18 and 5 non-overlapping clusters detected in Glu and MGE-GABA neurons, respectively (**Fig. S4E**). Four Glu clusters contained a human-specific DE gene (*OPRM1, GULP1, NIT2, PKDCC*). Several genes with important roles in the nervous system or neuropsychiatric disorders overlapped with the human up-DA GRE clusters. In particular, *CDH8* encodes a neurite outgrowth-regulating membrane protein cadherin-8 and has been linked to ASD and learning disability (25). Likewise, *ASTN2* is located within a Glu cluster on chromosome 9 and has been associated with ASD (27). Among the 76 genes located at Glu-neuronal DA clusters, 9 are associated with ASD according to the SFARI database (https://www.sfari.org/resource/sfari-gene/) (54), which is a significant enrichment (p = 0.02 by hypergeometric distribution). A previously described class of human genomic regions that harbor signatures of human-specific evolutionary changes consists of human-accelerated regions (HARs) (55). Notably, three of the 18 Glu clusters overlapped with HARs (HAR2, HAR22, HAR47; Fisher’s exact test, p = 0.005,), and each of these 3 HARs overlapped within a human Glu enhancer identified in our study.

We were particularly interested in the cluster containing *OPRM1*, a gene that encodes a receptor of endogenous opioid peptides and is implicated in opioid addiction (**Figs. 4B-E**) (18, 19). The *OPRM1* locus overlaps with one of the Glu clusters on chromosome 6 that contains 13 human- and Glu-specific up-DA enhancers as well as a human- and Glu-specific up-DA promoter (**Fig. 4B**). Concordant with these regulatory changes, *OPRM1* expression is upregulated in human compared with the two primate species in Glu but not in MGE-GABA neurons (**Fig. 4E**). We also found that two human-specific up-DA GREs within *OPRM1* harbor risk SNPs associated with opioid addiction (*rs1799971* and *rs3778150)*, which are situated within the *OPRM1* promoter and one of its enhancers, respectively (56, 57) (**Fig. 4C**). Strikingly, the major allele nucleotide of *rs3778150* (T>C) is a strictly human-specific substitution within a region which otherwise shows high sequence conservation across mammals (**Fig. 4D**). The *OPRM1* regulatory domain also provides an example of a respecification event, as it contains an enhancer that is active in Glu neurons in human but in MGE-GABA neurons in rhesus macaque (**Fig. 4B**). Thus, the *OPRM1* locus shows strong evidence of extensive evolutionary changes of its regulatory landscape that is coupled with concordant changes in gene expression specific to Glu neurons. In addition to *OPRM1*, two other genes implicated in drug abuse, *SLC17A8* (discussed above, **Fig. 3J**) and *PENK* (**Fig. S4F, G**), showed concordant evolutionary changes in gene expression and regulation. In contrast to *OPRM1*, the changes in these two genes were specific to MGE-GABA neurons. These findings suggest human-specific modification of cellular and molecular pathways implicated in drug addiction.

## DISCUSSION

Here, we demonstrate the value of analyzing isolated neuronal cell populations to increase the sensitivity and specificity of gene expression and GRE analysis compared with analyses of bulk cortical tissue. Our findings provide insight into the cell-type-specific regulatory landscape of the primate brain and its evolution. Our analyses show that neuron-subtype-specific regulatory elements preferentially changed during primate evolution as compared with elements that are active in multiple neuronal subtypes. This greater evolutionary plasiticity of neuron-subtype-specific GRE suggests that an alteration of a regulatory element that is beneficial or neutral when it occurs in a single cell type could be detrimental when it involves multiple cell and/or tissue types (58). We also detected regulatory elements that, despite being active in both Glu and MGE-GABA neurons of one species, were active in a specific cell type in other species. Thus, the assumption that a GRE that is active in multiple cell types (or tissues) represents a functionally conserved regulatory element is likely to be overly simplistic. Rather, these enhancer regions might employ different sets of transcription factor binding sites in different cell types and species, activating distinct regulatory programs.

Supporting these observations, we identified numerous regulatory elements whose activity showed neuron-subtype-specific evolutionary changes in humans (5,259 in Glu and 2,415 in MGE-GABA). We also detected 165 genes in Glu and 80 genes in MGE-GABA in humans with strong evidence of concordant evolutionary changes in their expression and the activity of at least one GRE. For the majority of these genes, the evolutionary change was detected across the entire regulatory domain (quantified as the number of evolved enhancers per gene and/or the magnitude of the enhancer H3K27ac signal). These findings demonstrate the functional importance of regulatory evolution in different neuronal subtypes. For instance, enrichment of H3K27ac in the promoter and enhancers of the catalase gene (*CAT)* in human Glu neurons is associated with a concordant human-specific upregulation of gene expression, which could protect these neurons against oxidative stress caused by the high metabolic activity of the human brain (42-44). The limitation of our study is that genes were linked to their putative enhancers using the distance-based assignment, which is complicated by the mostly unknown high-order chromatin structure. The latter can be assessed by genome-wide chromosome conformation capture (Hi-C) (59, 60). However, currently, Hi-C data are not available for Glu or MGE-GABA subtypes.

Complex spoken language is unique for humans, and the development of language was essential to human evolution, through enabling efficient exchange of information, higher order social organization as well as specializations of cognition and symbolic thinking. Our study emphasizes the importance of regulatory evolution in the development of language abilities and in the emergence of disorders associated with language. We identified two language-associated genes with strong human-specific (*ATP2C2*) or hominid-specific (*DCDC2*) as well as neuron-subtype-dependent concordant changes in expression and regulatory landscapes. We also discovered human-specific regulatory changes in Glu and/or GABA neurons in the *FOXP2* and *CNTNAP2* genes that are crucial for brain development, neural plasticity and language abilities (2, 23). *FOXP2* encodes Forkhead box protein P2 (FoxP2), a transcription factor that is expressed at high levels in the brain during fetal development (61). Many FoxP2 targets (including *CNTNAP2*) are implicated in schizophrenia and ASD (62, 63). ASD is characterized by difficulties in social communication (including language), and previous analyses have uncovered a link between ASD and HARs, suggesting that certain aspects of ASD evolved specifically in humans (64). Notably, here, we found that human-specific regulatory elements in neurons are not uniformly distributed in the human genome, and that clusters of these GREs in Glu neurons are enriched for HARs and genes associated with ASD.

Unexpectedly, we also identified concordant evolutionary changes in GREs and expression for several genes that have been implicated in drug addiction, with *OPRM1* showing an extensive re-organization of its regulatory domain and a pronounced upregulation of expression, specifically, in human Glu neurons compared with chimpanzee and rhesus macaque. Because *OPRM1* is one of the strongest candidates for affecting risk for opioid use disorder (56, 57), our findings emphasize that vulnerability to opioid addiction might have a unique human component (65).

The findings presented in this work highlight the importance of differential regulatory changes in major neuronal subtypes in brain evolution and brain disorders. These results call for further analyses that will connect evolutionary changes in regulation with those in gene expression in multiple subpopulations of neuronal and glial cells, in different brain areas, and across developmental trajectories, perhaps, using single-cell based approaches currently under development (14). These future studies will help to uncover the complex regulatory interaction networks that underlie the evolution of human brain function and human-specific traits, and hence will further advance the understanding of neuropsychiatric disorders.

## METHODS

### Specimens

Human brain specimens (DLPFC tissue samples from 4 clinically unremarkable male subjects) were described in (17) (**Table S1**). Tissue samples were dissected from the lateral part of Brodmann area 9 (BA9). The rhesus macaque brain tissues were obtained from the Texas Biomedical Research Institute and California National Primate Research Center from their established biospecimen distribution programs (**Table S1**). The chimpanzee brains were obtained from the National Chimpanzee Brain Resource. Brains of both rhesus macaques and chimpanzees were collected and snap frozen at the time of necropsy within 5 hours or less postmortem. The DLPFC was sampled from frozen brains in a region corresponding to BA9 as described in macaques and chimpanzees (66).

### Glu and MGE-GABA nuclei isolation by FANS

The isolation of nuclei was performed by FANS as described in (17). In short, to distinguish between the two neuronal populations, we employed antibodies against RNA-Binding Protein RBFOX3 (aka NeuN; mouse anti-NeuN, Alexa488-conjugated, MAB377X, Millipore) that is expressed in all neuronal nuclei, and antibodies against SOX6 (guinea pig anti-SOX6) (67). SOX6 is a transcription factor that regulates the ontogeny of the MGE-derived GABA neurons and is robustly expressed in these cells in the adult human PFC. This experimental approach has been previously validated using immunocytochemistry, immunohistochemistry and RNA-seq (15). As we previously reported (15), in addition to glutamatergic neurons, the FANS-isolated Glu population contained a small fraction of non-MGE GABA neurons (∼8% of the all sorted Glu neurons) (15). We used ∼800 mg and 150 mg of primate tissue to isolate nuclei for ChIP-seq and RNA-seq, respectively.

### ChIP-seq

H3K27ac ChIP was performed for each subject and neuronal subtype as described in (16, 17), using 100,000-150,000 FANS-separated nuclei and anti-H3K27ac antibody (rabbit polyclonal, Active Motif cat# 39133). We employed the “native” ChIP protocol (N-ChIP) that uses enzymatic chromatin fragmentation with micrococcal nuclease (MNase) without cross-linking proteins to DNA (16, 68). This approach yields ChIP-Seq data with a high signal-to-noise ratio and is, therefore, advantageous for profiling various histone modifications. ChIP-Seq libraries were constructed using NEBNext Ultra DNA Library Prep kit (New England Biolabs) as described in (17). For each sample, both ChIP and input (MNase-digested chromatin) libraries were generated and sequenced. Sequencing was performed on an Illumina HiSeq 2500 instrument, using a paired-end 50 (PE50) protocol to an average of ∼40 million read pairs per sample.

### RNA-seq

RNA was isolated from each subject and neuronal subtype as described (17), using 40,000 FANS-separated nuclei. To preserve RNA integrity, the RNAse inhibitor (Clontech) was added during the each step of the nuclear preparation. Nuclei were sorted directly into tubes containing 3:1 by volume of the Extraction Buffer from PicoPure RNA Isolation kit (ThermoFisher Scientific). RNA was then extracted from the sorted nuclei using the PicoPure kit, and the RNA-seq libraries were constructed using SMARTer Stranded Total Pico-Input RNA-seq kit (Clontech) and 10 ng RNA from each sample, as described (17). Libraries were sequenced on HiSeq 2500, using PE50 protocol to an average of ∼50 million read pairs per sample.

### Data analysis methods

see **Supplementary Methods**

## ACKNOWLEDGMENTS

This work was supported by PsychEncode consortium NIH/NIMH MH103877 and MH122590 (S.D.), NIH/NIDA DA043247 (S.D.), VA Merit Awards BX001829 and BX002876 (S.D.), Intramural funds of the U.S. Department of Health and Human Services to National Library of Medicine (E.V.K.), Dutch Royal Academy of Sciences (KNAW; M.P.C.); Parkinson’s foundation (Stichting ParkinsonFonds, M.P.C.); James S. McDonnell Foundation 220020293 (C.C.S.); NSF INSPIRE SMA-1542848 (C.C.S.). The chimpanzee brains were obtained from the National Chimpanzee Brain Resource (supported by NIH/NINDS NS092988). We gratefully acknowledge the help of Cheryl Stimpson for technical assistance with chimpanzee brain dissections and the help of Dr. Rob Patro with the use of Salmon software package.

