## Supplementary figures and methods for "Evolution of regulatory signatures in primate cortical neurons at cell type resolution"

**SUPPLEMENTARY METHODS**

**RNA-seq data analysis:** The comparison of gene expression between primate species is complicated by the inferior quality of the available gene annotations for chimpanzee and rhesus macaque as compared to human. To overcome this difficulty, we generated improved transcriptome annotations for chimpanzee and rhesus macaque, using previously published high-coverage RNA-seq datasets for bulk brain tissue from these two species (1, 2). The RNA-seq datasets were mapped by STAR [version 2.5, **github.com/alexdobin/STAR**] to the rheMac8 or panTro5 genome builds. Next, transcriptome annotations were generated using StringTie [version 1.3, **ccb.jhu.edu/software/stringtie**] with default parameters, assembled into consensus annotations using TACO [version 0.7.3, **tacorna.github.io**], and merged together with the Ensembl v92 annotations using Cuffmerge [version 2.2.1, **cole-trapnell-lab.github.io/cufflinks/cuffmerge**]. The resulting transcriptome annotations contained ~40% more unique exons (~360,000-400,000) compared to the chimpanzee and rhesus macaque annotations that are available from Ensembl.

To limit potential biases in gene expression quantification across primate species further, such as the differences in transcript size, we adopted an approach reported in (3). For each species and gene, we constructed the sets of "metaexons" by merging together the overlapping exons. We then defined "orthologous metaexons" as those metaexons that had unique reciprocal equivalents in all three species. We used the UCSC LiftOver tool with default parameters convert coordinates between species. Next, for each species, we filtered the transcriptome annotations (generated as described above) by keeping only those transcript segments which comprised contiguous sets of orthologous metaexons. Single-exon transcripts which belonged to multi-exon genes were excluded to avoid spurious annotation entries. Because the lengths of 3-UTR transcript sequences varied significantly between species, the 3-UTR sequences were trimmed. The resulting curated annotations (in GTF format) covered 163,650 orthologous metaexons and 22,898 genes (including 16,847 protein-coding genes). We then used these GTF files to generate FASTA transcriptome files for each species using RSEM [version 1.2.25, **github.com/deweylab/RSEM**].

RNA-seq data sets (FASTQ files) generated in the current study for the three species and two neuronal subtypes were pre-processed by trimming the adapters using the Scythe software tool (**github.com/vsbuffalo/scythe**) and by removing low quality reads using Sickle (**github.com/najoshi/sickle**). Data files were further trimmed by 3 bp from the 5′ end of Read 1, in accordance with the SMARTer RNA-seq library kit protocol, using seqtk (**github.com/lh3/seqtk**). The pre-processed FASTQ files were then analyzed together with the newly generated curated annotations (in FASTA transcriptome format) using a transcript-quantification software tool Salmon [version 0.9.1, **combine-lab.github.io/salmon**]. The FASTA transcriptome files were pre-converted to Salmon indexes using a Salmon command "*index*" with the k-mer minimum length setting *"-k 23*". GC%-bias correction option was included during the Salmon quantification step. The Salmon command line was as follows: *salmon quant -I $INDEX -l A –incompatPrior 0.0 –gcBias -1 $SEQ_READ1 -2 $SEQ_READ2 -p 16 -o $TRANSCRIPTS_QUANT*. The output of the Salmon command was aggregated by gene using the R package "tximport" [version 1.6.0] and then further analyzed for differential expression by the R package "DESeq2" [version 1.18.1, (4)]. The genes with low read counts were removed with the "independent filtering" option during the DESeq2 analysis. Differentially expressed (DE) genes were defined using the settings: abs(fold change) > 2, FDR < 0.05. Only protein-coding and lincRNA genes were retained for the subsequent analyses.

**ChIP-seq data analysis**

**1. GRE identification in human, chimpanzee and rhesus macaque:** Paired-end sequences were aligned using Bowtie [version 1.1.2, **bowtie-bio.sourceforge.net**] with the following settings: *chunkmbs 512 -S -q -l 40 -n 1 -phred33-quals -p 7 -m 1*, therefore excluding reads with more than one mismatch or with multiple alignments. Mapping was done onto rheMac8 for rhesus macaque, panTro5 for chimpanzee, and hg38 for human. Between 28 million and 68 million reads were uniquely mapped for each of the samples, with an average mapping percentage of 81% (**Table S2**). FriP scores ranged from 27% to 50%, thus largely exceeding the 1% threshold used by ENCODE (5). Significant H3K27ac-enriched regions (peaks) were called using MACS2 version 2.1.1.20160309 using an input control (MNase-digested chromatin) for each sample with the following settings: *callpeak -B -f BAMPE –broad –broad-cutoff 0.00001*. Peaks that were called in three out of four replicates were maintained, merged when overlapping, and the resulting peaks smaller than 2,000 base pairs (bp) were extended to a minimum size of 2,000 bp (peak center ± 1,000 bp) as described in (6). Total (non-redundant) sets of 78,086 and 65,527 (in human), 75,156 and 62,378 (in chimpanzee), and 82,347 and 72,457 (in rhesus macaque) H3K27ac-enriched regions (peaks) were compiled for Glu and MGE-GABA, respectively (**Table S3**). To perform differential enrichment analysis between the two cell types (**Fig. 1E** and **Fig. S1E**), H3K27ac-enriched regions (putative GREs) in Glu and MGE-GABA neurons were merged within each species, resulting in 102,080 GREs for human, 92,613 GREs for chimpanzee, and 106,883 GREs for rhesus macaque.

**2. Cross-species alignment of GREs and differential acetylation analysis:** Coordinates for GRE comparison between the 3 primate genomes were obtained as previously described (6). In brief, hg38 coordinates were acquired for Glu and MGE-GABA H3K27ac-enriched regions in chimpanzee or rhesus macaque using the UCSC LiftOver tool (minMatch = 0.5). Because mapping onto the target genome and back to the source genome (reciprocal liftOver) had to be unique, the regions that changed more than 50% in size by the liftOver procedure were discarded. For chimpanzee, 74985 out of 75156 Glu GREs and 62245 out of 62378 MGE-GABA GREs were mapped onto hg38. For rhesus macaque, 81183 out of 82347 Glu GREs and 71628 out of 72457 MGE-GABA GREs were mapped onto hg38.

For pairwise comparison between human and rhesus macaque, in each neuronal subtype, the coordinates for rhesus macaque GREs were mapped to hg38 by liftOver and merged with human GREs. Similarly, for the comparison across the three primate species, the hg38 coordinates for chimpanzee and rhesus macaque GREs were merged with the human GREs. As merging of (partially) overlapping regions of the three species results in changed coordinates, these regions were then subjected to the reciprocal liftOver conversion from hg38 back to panTro5 or rheMac8 genomes in order to obtain three-way orthologues. The regions that changed more than 50% in size by the liftOver procedure as well as GREs with zero reads in all replicates in each species were discarded. We applied the following criteria to select the enriched regions with orthologues on all 3 genomes in order to correct for genome quality differences between the species: *(1)* ≥ 90% of the bases had to be known in each of the reference genomes (<10% overlap with UCSC Table Browser's Gap Locations lists), and *(2)* a peak signal was not allowed to change significantly if multi-mapping of reads was allowed; such peaks could contain duplicated or repetitive regions that are more susceptible to poor annotation in the lower-quality nonhuman primate genomes. To identify these "MultiMap" regions, reads for each sample were mapped with the following settings using bowtie: *–best–strata -M 1*. The called peaks were discarded if they were not significantly enriched using unique mapping settings. The peaks that did not satisfy criteria *(1)* and *(2)* were "blacklisted" and excluded from the analysis. In addition, we excluded the GREs which were located at a distance >1 Mb from the nearest TSS and thus could not be annotated to a gene using GREAT (981 in Glu and 512 in MGE-GABA neurons, respectively). Following this strategy, we obtained 99,339 Glu and 82,761 MGE-GABA GREs in the comparison between human, chimpanzee, and rhesus macaque, and 93,222 Glu and 79,181 MGE-GABA GREs in the comparison between human and rhesus macaque (**Table S6**). In order to perform Pearson correlation analysis, hierarchical clustering and PCA on ChIP-seq data from 3 species and 2 cell types simultaneously, the above-generated Glu and MGE-GABA GREs were merged on the hg38 genome and subjected to reciprocal liftover and filtering as described above, resulting in a data set of 122,462 GREs (**Table S20**).

Differentially acetylated (DA) GREs were detected with DESeq2 (4) by pairwise comparison of the four independent biological replicates between cell types or species, using the following criteria: FDR < 0.05, Log2 FC > 2. Read counts were normalized for visualization and regression using the rlog function of DESeq2, which transforms the count data to the log2 scale in a way which minimizes differences between samples for rows with small counts and normalizes the data with respect to a library size. All GREs were used for comparisons. This was followed by splitting GREs into enhancers and promoters.

**GRE-to-gene annotation and gene ontology analysis** of cell type- or species-specific GREs was performed using GREAT (**http://great.stanford.edu**). Gene regulatory domain was defined as following: each gene was assigned a basal regulatory domain of a 1 kb distance upstream and downstream of the TSS (regardless of other nearby genes). The gene regulatory domain was extended in both directions to the nearest gene's basal domain but no more than 1 Mb in one direction.

**Detection of clusters of human-specific upregulated GREs**: Human genome was subdivided into sets of non-overlapping, equally-sized 1 Mb bins, with each set shifted by 100 kb. For each bin, the number and positions of human-specific (vs. chimpanzees and rhesus macaques) upregulated GREs were recorded. The analysis was performed separately for Glu and MGE-GABA GREs, and differentially acetylated GREs were defined as those with FC > 2 at FDR < 0.01 between the neuronal subtypes for this analysis. The enrichment of human-specific GREs within each bin was assessed using two complementary approaches, and only the bins which were found significant by both analyses were considered significantly enriched.

In the first analysis, we modeled the random distribution of Glu- or MGE-GABA-neuronal human-specific GREs across the mappable fraction of the human genome (2.7e9 bp, as per MACS2 settings) using the Poisson distribution. After correction for multiple testing, the number of human-specific GREs per bin that was >11 for Glu GREs and >8 for MGE-GABA GREs corresponded to a significant enrichment (adjusted p-value < 0.05).

In the second analysis, we assessed if an increased H3K27ac signal within human-specific GREs in a particular bin could be explained by the increased total H3K27ac signal for all human GREs within that bin. To this end, human GREs were subdivided into human-specific GREs and non-DA GREs (human GREs which were not DA vs. chimpanzee or rhesus macaques). We counted the number of human-specific GREs and non-DA GREs within each 1 Mb bin using the data for the 4 human samples in Glu or MGE-GABA neurons. We then tested whether the human-specific GREs were more abundant than non-DA GREs using Poisson distribution, with mean given by the number of non-enriched GREs multiplied by a scaling factor to account for the ~10-fold lower total number of human-specific GREs. Only the bins with significant increase in the human-specific signal compared with the background (non-DA) signal were retained (FDR < 0.05). Finally, we combined the results obtained in the two analyses. Merging the overlapping bins located within the same genomic region resulted in 18 and 5 human-specific GRE clusters in Glu and MGE-GABA neurons, respectively.

**Analysis of correlation between inter-species and inter-cell-type changes in enhancer signal intensity**: We used a threshold-free approach to assess the correlation between changes in enhancer signal intensity (fold change (FC) of normalized H3K27ac read counts) in the interspecies comparison (Hu vs. Rh) compared with inter-cell-type comparison (Glu vs. MGE-GABA). The analysis was performed separately for enhancers detected in Glu or MGE-GABA. For example, for human Glu enhancers, we directly compared the neuronal subtype specificity (e.g., difference between Glu and MGE-GABA in human, $Log2\ cell\ type\ FC_{,Hu} = Log2\ (H3K27ac_{Glu,Hu}\ /\ H3K27ac_{GABA,Hu})$ with the evolutionary change (difference between human and rhesus macaque in Glu cells, $Log2\ species\ FC_{,Glu} = Log2\ (H3K27ac_{Glu,Hu}\ /\ H3K27ac_{Glu,Rh})$. This analysis could be affected by spurious correlations if the same datasets were used in multiple steps of analysis (double-dipping), such as when the empirically determined strength of each enhancer in human Glu cells, $H3K27ac_{Glu,Hu}$, was used to estimate both $Log2\ cell\ type\ FC_{,Hu}$ and $Log2\ species\ FC_{,Glu}$. To ensure a valid statistical comparison, we therefore randomly selected independent biological samples for determining enhancer regions (donors 1 and 2), for estimating the species difference (donor 3), and for estimating the cell type difference (donor 4). We first randomly selected two of the four human samples (donors 1 and 2), called H3K27ac peaks with MACS2 in Glu and MGE-GABA neurons using data for these two samples only, and then merged peaks that were detected in both replicates (intersect by at least 1bp). Peaks that had a length less than 2kb were extended to +/-1kb from the center. We then identified regions orthologous to these GREs in the rhesus macaque genome (rheMac8) using LiftOver and kept only those regions which could be lifted over back to the human hg38 genome with exactly the same coordinates. We further removed regions that had > 50% change in length after LiftOver and regions that have >10% overlap with genomic gaps in either species. The remaining regions were grouped into TSSs and enhancers, using the +/-1kb regions from TSSs annotated by RefSeq, resulting in 52,828 and 45,899 enhancers in Glu and MGE-GABA, respectively. Read counts within the peaks were calculated using featureCounts (R package "subread") and normalized using the median of ratios method in DESeq2. Fold changes were calculated using the two remaining replicate human samples and a randomly selected rhesus macaque sample, as described above. Shuffled controls were created by randomly reordering the values of fold changes between species and cell types. The differences between the observed and the shuffled data were estimated with a two-dimensional kernel density estimation in R.

**Detection of GREs with species-dependent and/or cell-type-dependent changes in H3K27ac signal intensity:** Glu- or MGE-GABA H3K27ac peaks (putative GREs) detected in human (Hu) and/or rhesus macaque (Rh) were combined (172,403 GREs in total) and the signal intensity of peaks in Glu and MGE-GABA neurons in each of the two species was estimated using rlog values in four replicates. Next, for each GRE, an ANOVA test was performed using the formula: *rlog ~ species + cell_type + species:cell_type* , where the interaction effect of species and cell type was accounted for by the third term. For each term in the ANOVA, the proportion of variance explained (eta squared, $\eta^2$) was used to estimate the effect size, and p-values were corrected for multiple comparisons using the Benjamini-Hochberg FDR method. Subsets of GREs with strong and significant effect detected by ANOVA were defined as follows: “species effect”: $\eta^2_{species} > 0.8$, $FDR_{species} < 0.05$; “cell type effect”: $\eta^2_{cell\_type} > 0.8$, $FDR_{cell\_type} < 0.05$; “interaction effect”: $\eta^2_{species:cell\_type} > 0.8$, $FDR_{species:cell\_type} < 0.05$; “mixed effect”: $\eta^2 < 0.5$, FDR > 0.05 for all three terms.

### SUPPLEMENTARY FIGURES

#### Figure S1 Regulatory changes in Glu and MGE-GABA neurons during primate evolution

**S1A.** Heatmaps showing Pearson correlation coefficients across H3K27ac ChIP-seq replicate samples for each condition. *Left column*: Glu samples; *right column*: MGE-GABA samples. *Top row*: human samples; *middle row*: chimpanzee samples; *bottom row*: rhesus macaque samples; four replicates for each species and cell type. The non-redundant lists of GREs that were identified in at least 3 out of 4 biological replicates for each species were used in the analysis. Hu, human; Ch, chimpanzee, Rh, rhesus macaque.

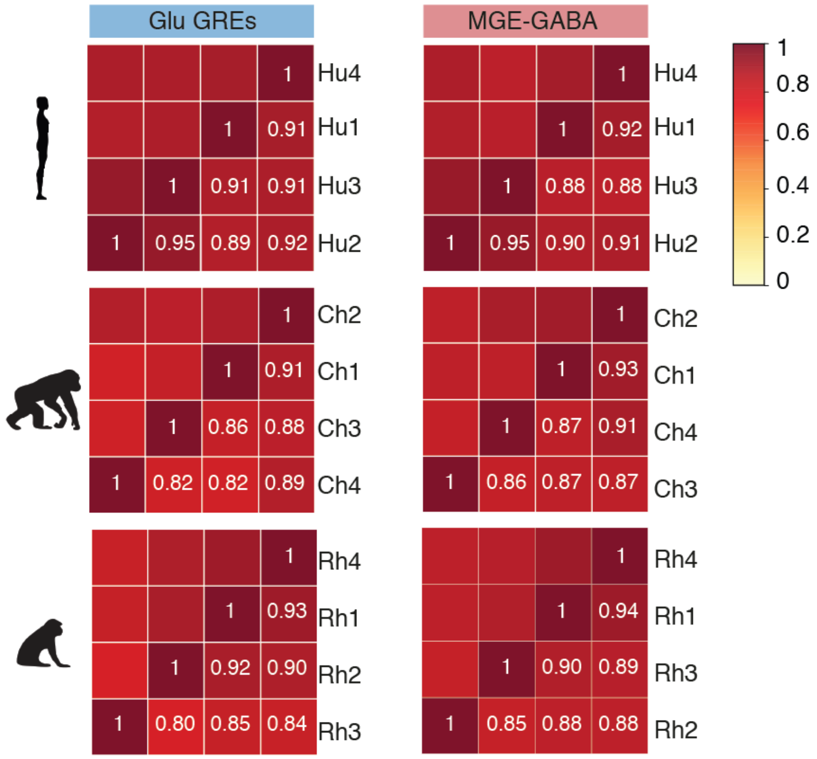

**S1B, C, D.** H3K27ac ChIP-seq profiles for human (**B**), chimpanzee (**C**), and rhesus macaque (**D**) in the vicinity of a Glu marker gene *TBR1* (*left panel* in each figure) and an MGE-GABA marker gene *SOX6* (*right panel* in each figure). Shown are read-per-million (RPM)-normalized ChIP-seq reads for H3K27ac (axis limit 3 RPM). The tracks represent four replicate samples for each species in Glu (*blue*) and MGE-GABA (*red*) neurons.

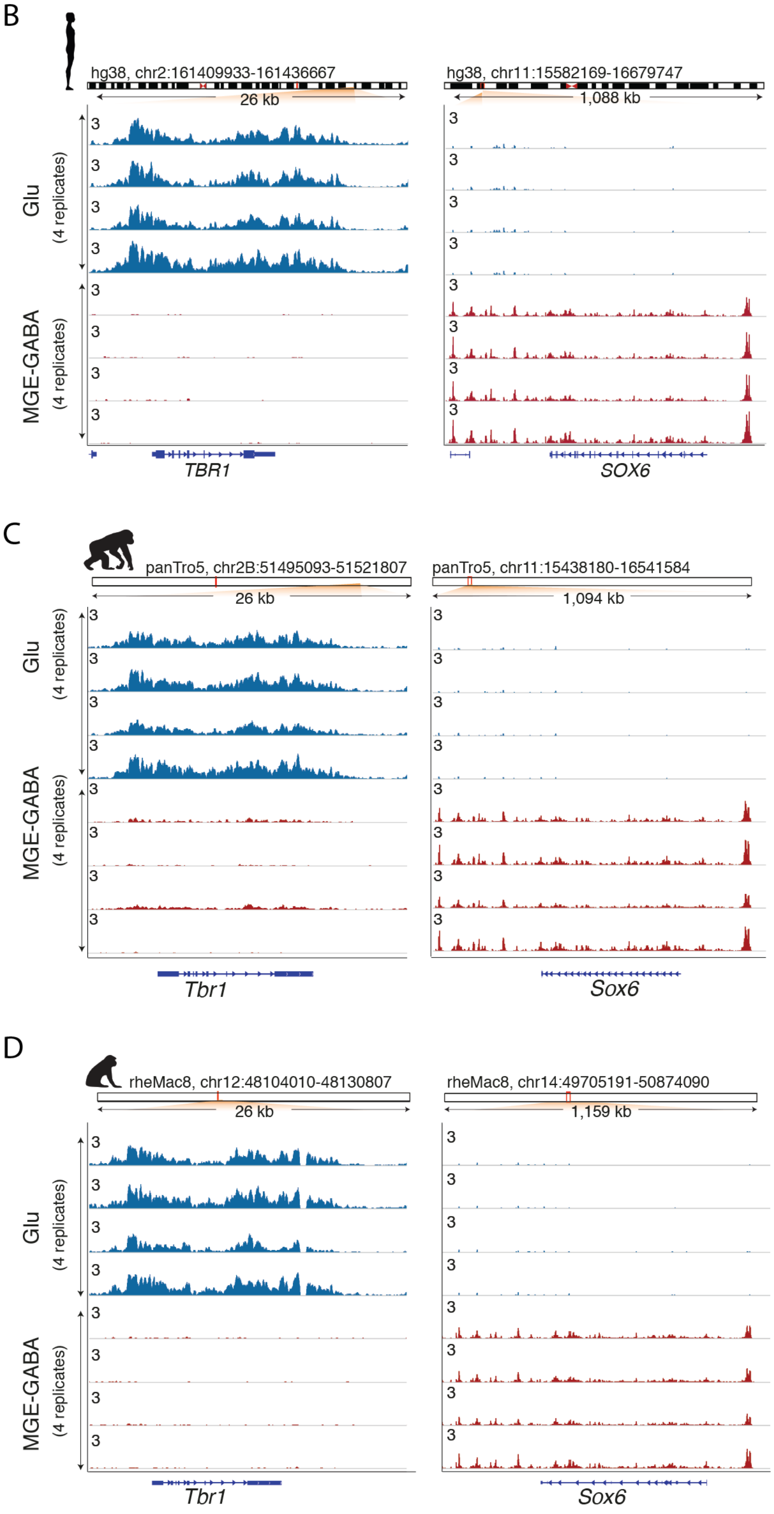

**S1E**. Fractions of GREs that are differentially acetylated (DA) or not differentially acetylated (non-DA) between Glu and MGE-GABA neurons in the 3 species. DA GREs were defined by DESeq2 (FC > 2, FDR < 0.05). The total number of GREs for each species is shown below the figure.

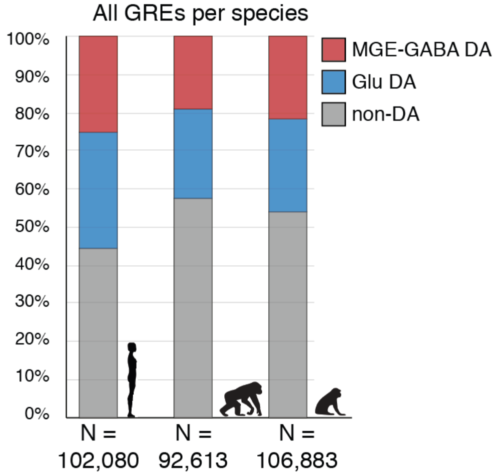

**S1F.** Principal component (PC) analysis of H3K27ac ChIP-seq data for Glu and MGE-GABA neurons human, chimpanzee and rhesus macaque. The non-redundant list of GREs from the 3 species and 2 neuronal subtypes was used for the analysis (**Methods**). Shown are the results for PC2 and PC3. See **Fig. 1H** for the ChIP-seq PCA results for PC1 and PC2.

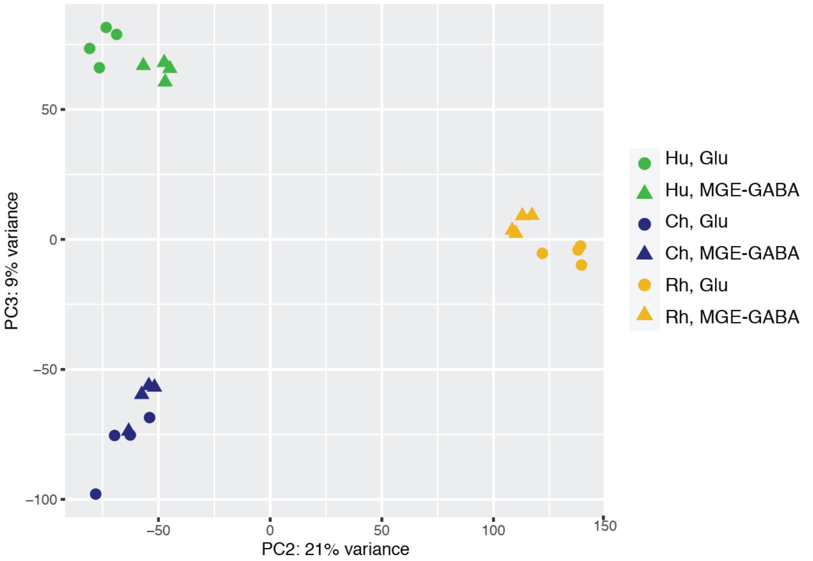


**S1G.** Hierarchical clustering heatmap showing Pearson correlation coefficients across Glu and MGE-GABA H3K27ac ChIP-seq samples from 3 species [human (Hu), chimpanzee (Ch) and rhesus macaque (Rh)]. The non-redundant list of GREs from the 3 species and 2 neuronal subtypes was used for the analysis (**Table S20**).

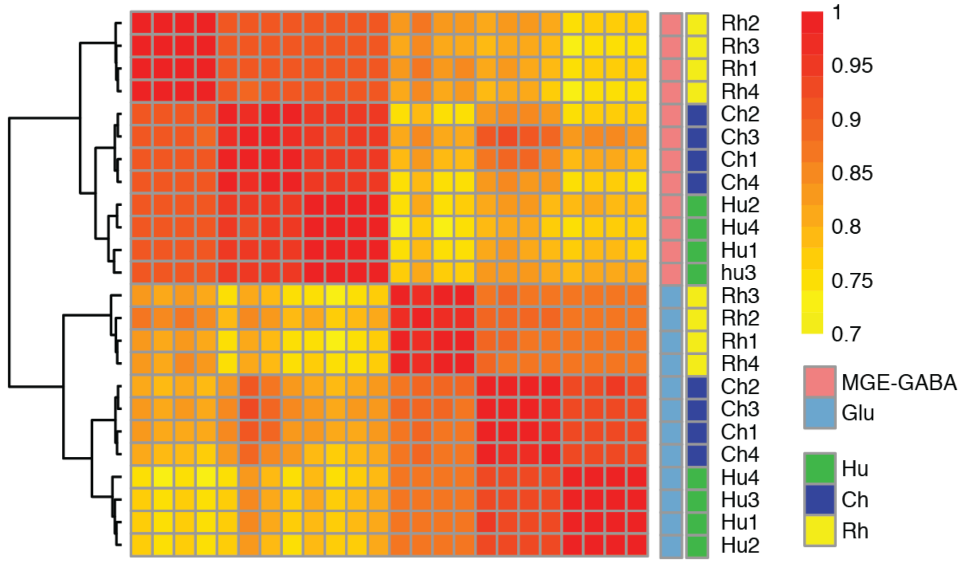

**S1H.** Numbers of differentially acetylated (DA) GREs detected in Glu or MGE-GABA neurons in pairwise comparisons of H3K27ac ChIP-seq datasets from Hu, Ch and Rh. The results are shown separately for enhancers (*left panel*) and promoters (*right panel*) GREs.

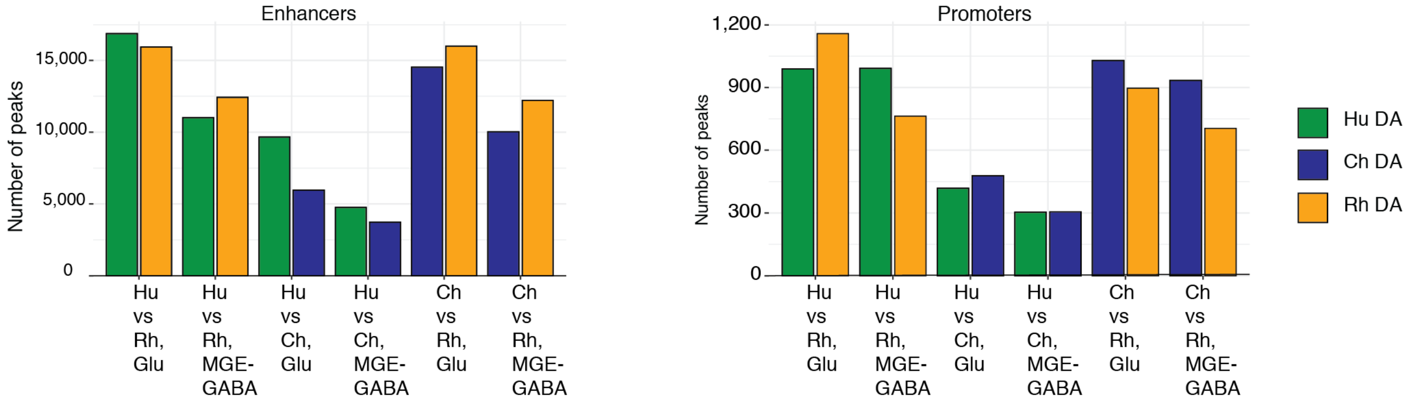

**Figure S2 Evolutionary changes in neuron-subtype-specific and pan-neuronal regulatory elements.**

**S2A.** Analysis of species specificity (Hu vs. Rh) of GREs that were differentially acetylated (DA) or non-differentially acetylated (non-DA) between Glu and MGE-GABA neurons. *Top panel:* Venn diagram showing the numbers of human GREs that were Glu DA (*blue*), MGE-GABA-DA (*red*) or non-DA (*purple*). *Bottom panel*: pie charts indicate the fractions of the GREs that are DA (*grey*) or non-DA (*purple*) between Hu and Rh. In both neuronal subtypes, neuron-subtype-specific GREs are more often DA between Hu and Rh than non-DA GREs (p values <2.2e-16 by Fisher's exact test).

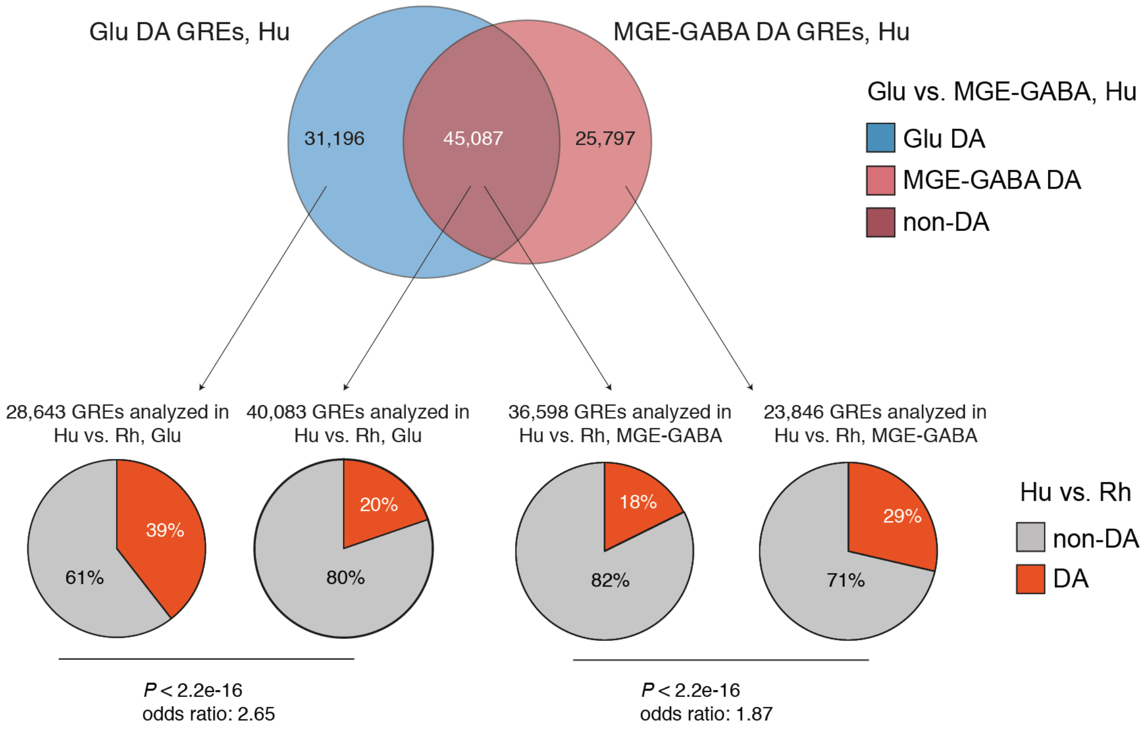

**S2B.** Verification of the findings depicted in **Fig. 2A** using a threshold-free approach. *Top row*: analysis for human Glu enhancers. *Bottom row*: analysis for human MGE-GABA enhancers. *Left columns*: observed data. Correlation between the fold changes (FC) in enhancer activities in the inter-species comparison (human vs. rhesus macaque) vs. inter-cell-type comparison (Glu vs. MGE-GABA) (see Methods). Each dot represents one enhancer. Spearman correlation coefficient and p-value are shown. *Middle columns*: shuffled data. Similar to the analysis in the left panel, but after randomly reordering the values of fold changes between species and cell types. *Right columns*: differences between the observed and shuffled data that were calculated using a two-dimensional kernel density estimation.

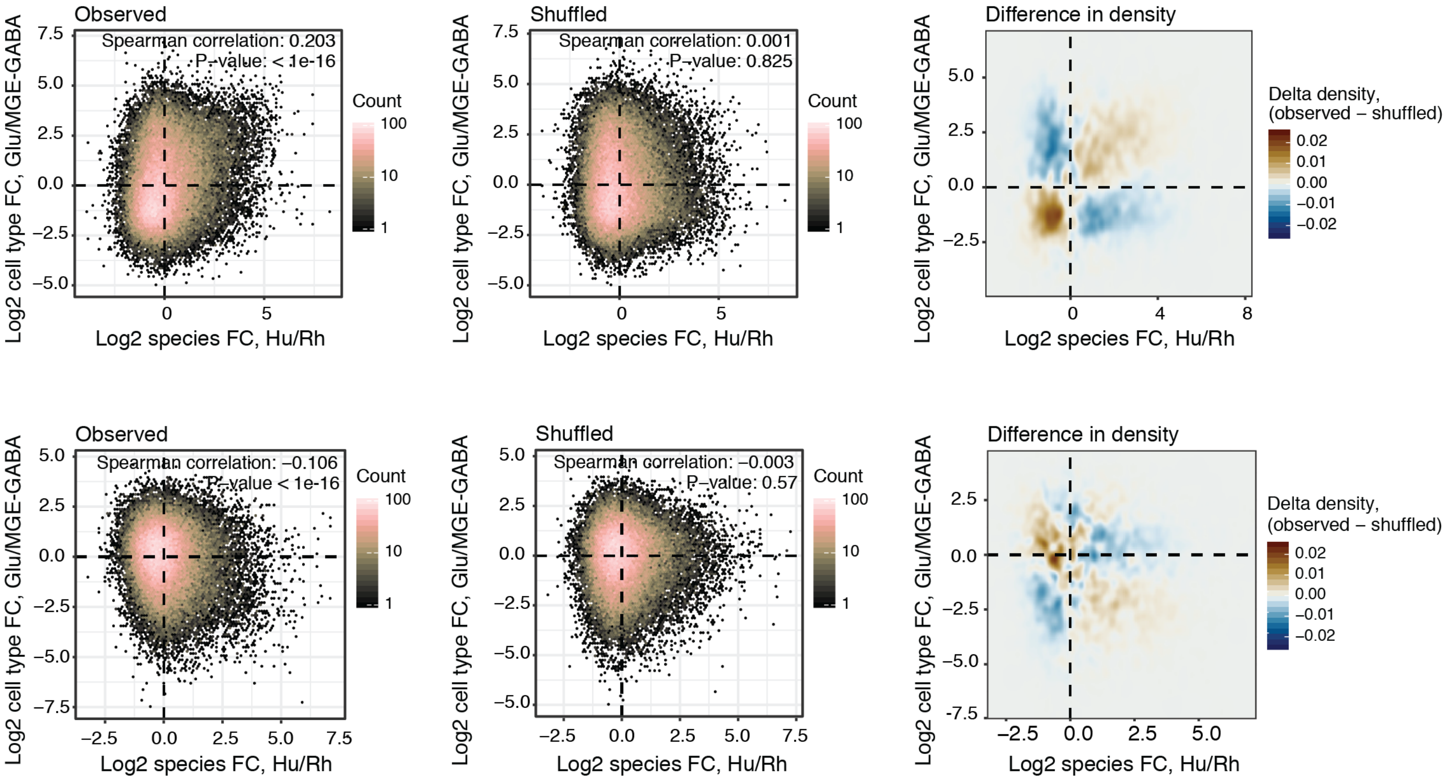

**S2C. Positionally conserved enhancers undergo neuron-subtype-specific evolutionary changes.**

Simultaneous cross-species and cross-cell-type comparisons of H3K27ac signal intensities for gene regulatory elements (GREs). The analysis included the GREs detected in Glu and/or MGE-GABA neurons in humans (Hu) and/or rhesus macaques (Rh). *Left panel*: evolutionary fold changes of H3K27ac signal intensities (Hu/Rh) were compared between Glu and MGE-GABA neuronal subtypes; *right panel*: neuronal-subtype-dependent fold changes (Glu/MGE-GABA) of H3K27ac signal intensities were compared between Hu and Rh. An ANOVA test was performed using the following formula:

*rlog ~ species + cell_type + species:cell_type*

Eta squared ($\eta^2$) was used as a metric to estimate effect size of each term in ANOVA; p-values were corrected for multiple comparisons using the Benjamini and Hochberg's FDR method. GREs with strong and significant effect detected by ANOVA ($\eta^2 > 0.8$, FDR < 0.05; **see Methods**) are highlighted in different colors. *Orange color*: significant evolutionary change in both cell types (10,307 GREs, "species effect" in ANOVA); *green color*: significant neuronal-subtype-dependent change (39,851 GREs, "cell type effect" in ANOVA); *purple color*: significant evolutionary respecification change (334 enhancers, "interaction effect" in ANOVA); *yellow color*: significant evolutionary change in one cell type only (12,657 GREs, "mixed effect"). By overlapping the results of this analysis with the DA analysis (**Fig. 2C**), we identified 3,074 enhancers that changed in the same direction in both cell types, 2,425 enhancers that changed in one but not in the other cell type, and 100 enhancers that underwent a respecification (**Fig. 2D, Table S9**).

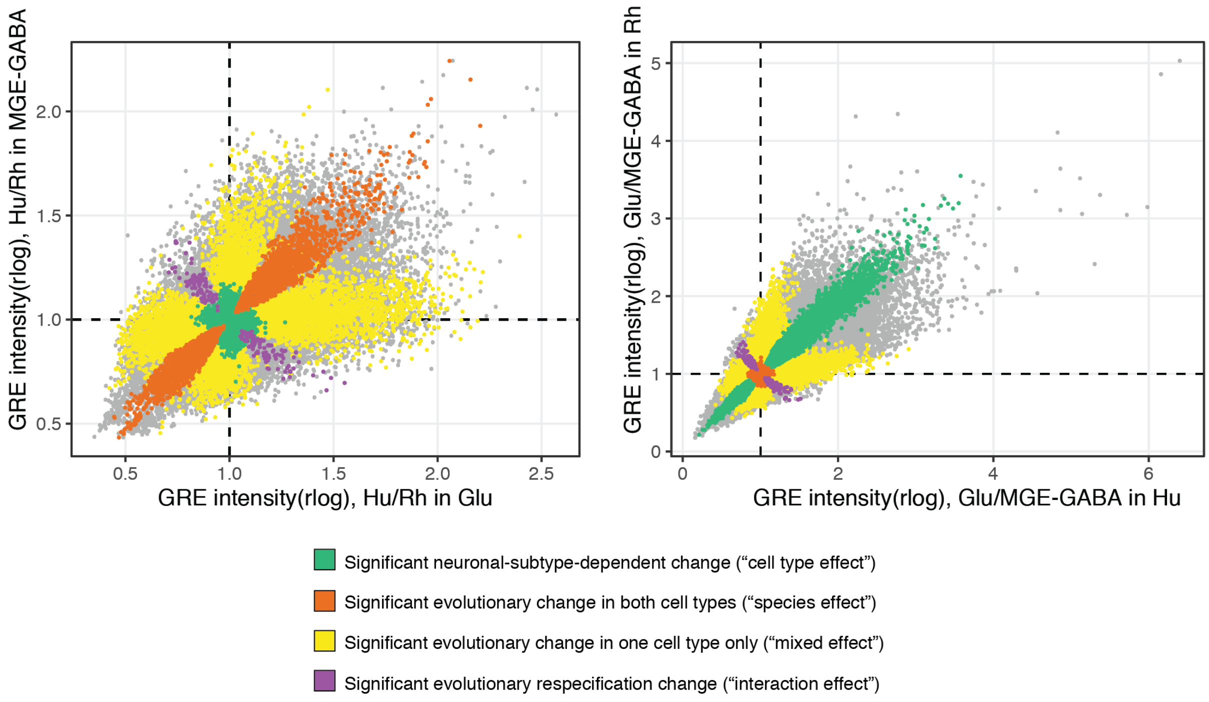

**Figure S3 Concordant evolutionary changes in GREs and gene expression.**

**S3A.** Heatmaps showing Pearson correlation coefficients across RNA-seq replicate samples for each condition (3 species and 2 neuronal subtypes in each species). Protein-coding and lincRNA genes were used in the analysis.

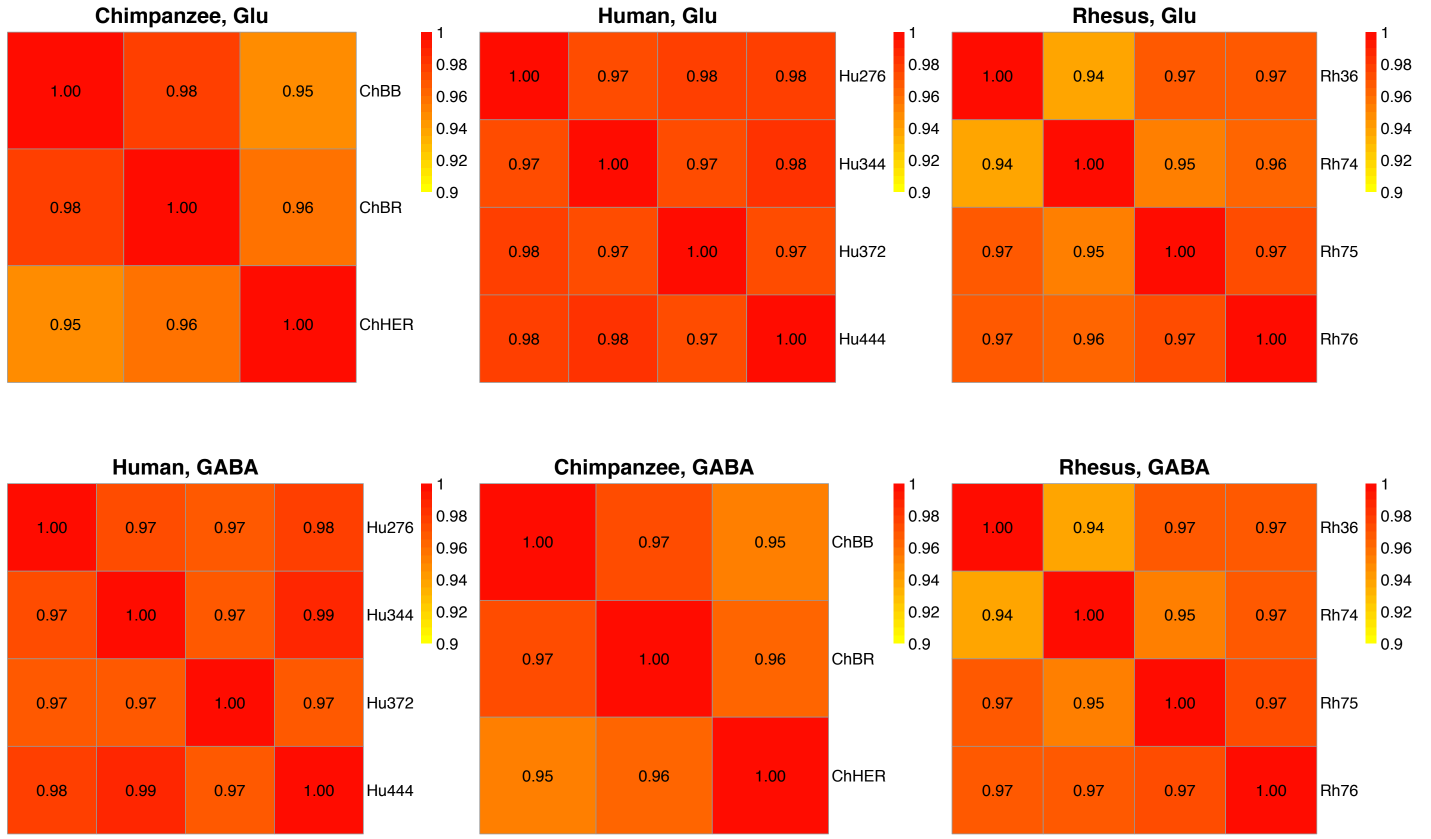

**S3B.** Pairwise comparison of gene expression between Glu and MGE-GABA neurons in each of the 3 species; *left panel*: human*; middle panel*: chimpanzee; *right panel*: rhesus macaque. Each dot represents one gene. TPM values are shown in the scatter plots, and differentially expressed genes called by DESeq2 (FDR<0.05) with at least 5-fold enrichment are highlighted. *Orange dots*: genes that show MGE-GABA-specific differential expression; *cyan dots*: genes that show Glu-specific differential expression; *dark blue dots*: genes that do not show neuron-subtype-specific differential expression. Examples of marker genes in each cell type are labelled. The results confirm the neuron-subtype-specificity of FANS-separated DLPFC Glu- and MGE-GABA-neuronal nuclei in the three species. Known specific markers of Glu neurons (*SLC17A6*, *SLC17A7*, *BDNF*, *SATB2*, *TYRO3*) or MGE-GABA neurons (*GAD2*, *SLC32A1*, *SOX6*, *LHX6*, *SST*, *PVALB*) show upregulated expression in FANS-separated Glu- or MGE-GABA- neuronal nuclei, respectively.

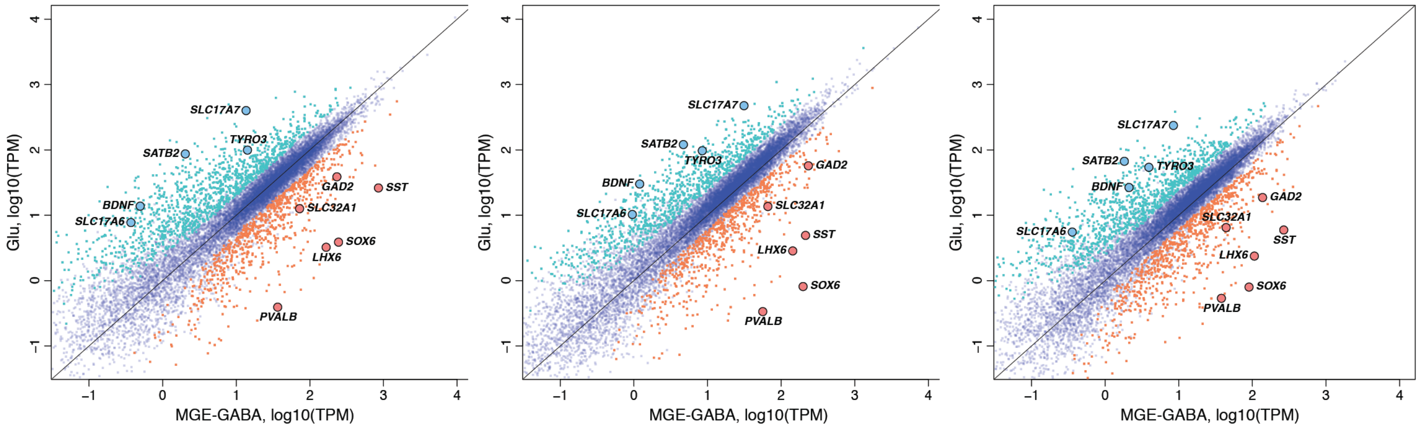

**S3C.** Hierarchical clustering heatmap showing Pearson correlation coefficients across Glu and MGE-GABA RNA-seq samples from human, chimpanzee and rhesus macaque subjects. Pearson correlation analysis was performed using regularized log2-transformed RNA-seq signal (rlog values). Protein-coding and lincRNA genes were included in the analysis.

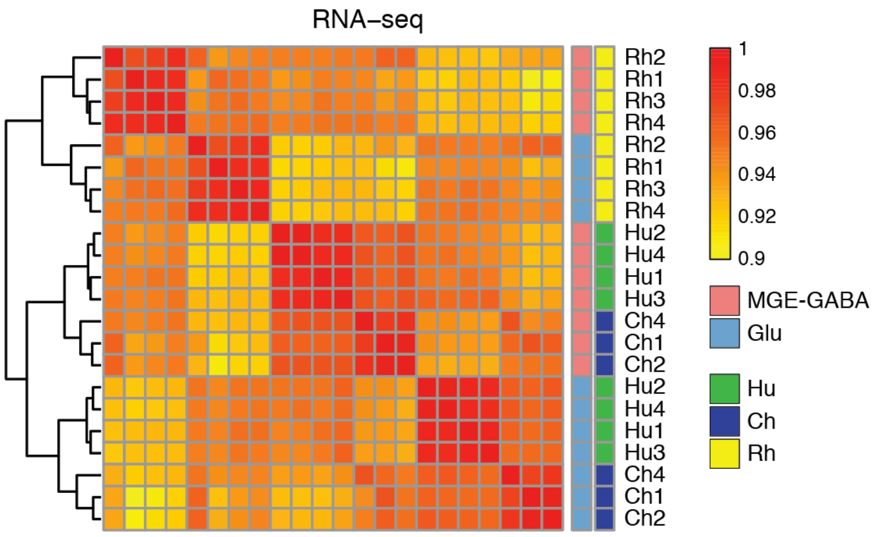


**S3D.** Principal component (PC) analysis of the RNA-seq data sets from (C). Shown are the results for PC1 and PC2.

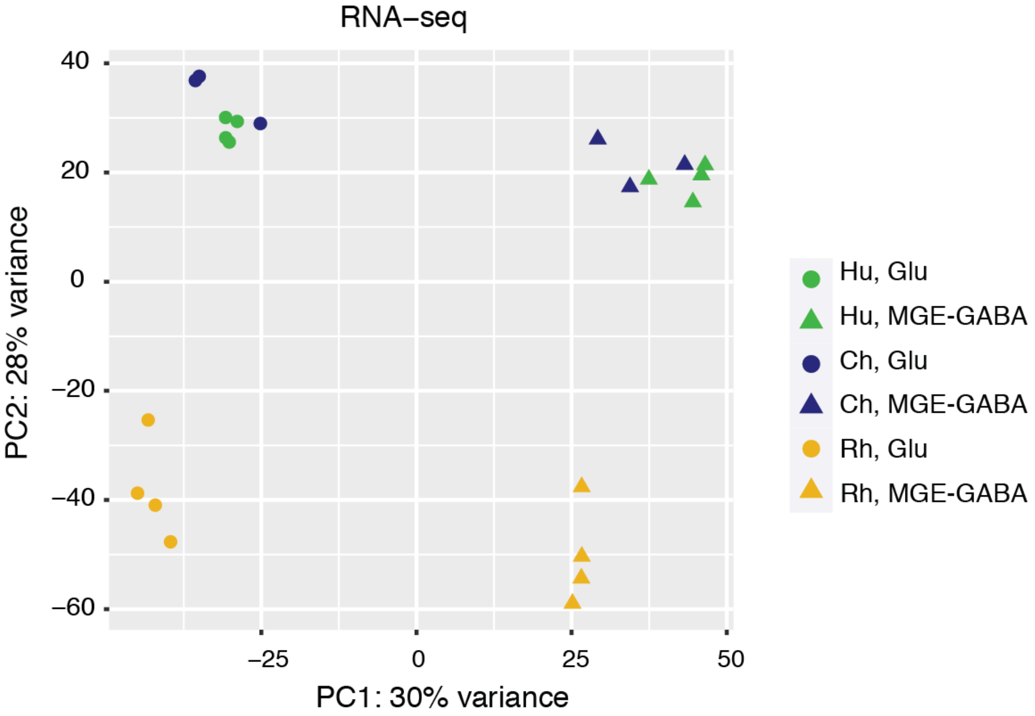

**S3E.** PC analysis of the RNA-seq data sets from (C). Shown are the results for PC2 and PC3.

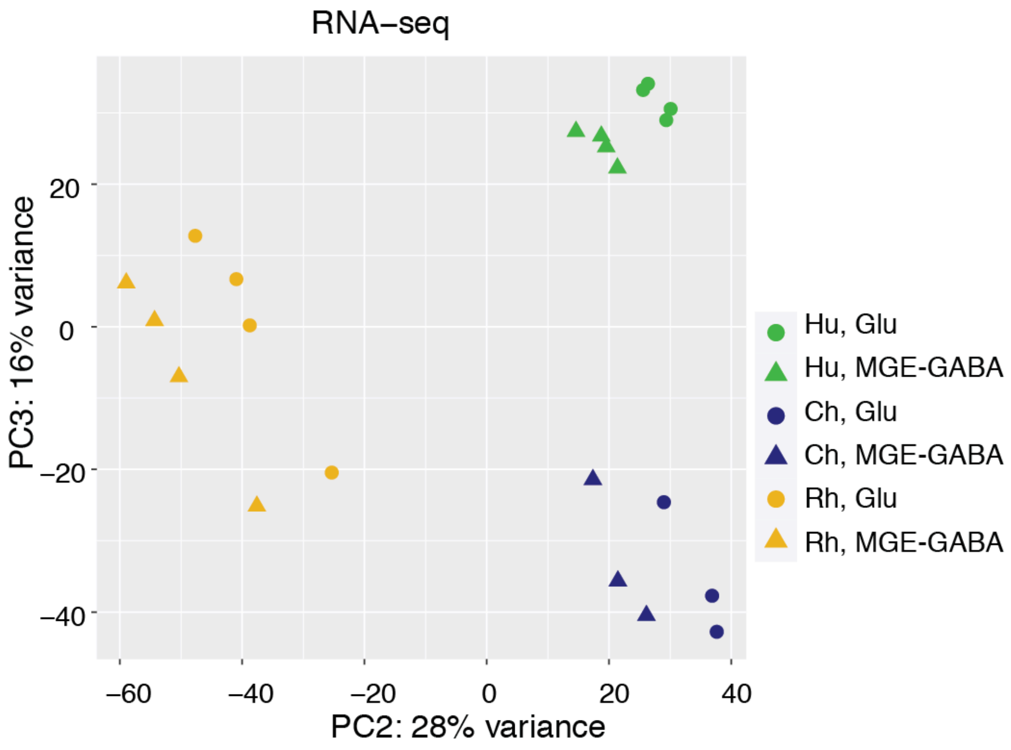


**S3F.** Evolutionary conservation of gene expression profiles for protein-coding and lincRNA genes. Shown are Spearman correlation coefficients for pairwise comparisons of RNA-seq datasets among human (Hu), chimpanzee (Ch) or rhesus macaque (Rh).

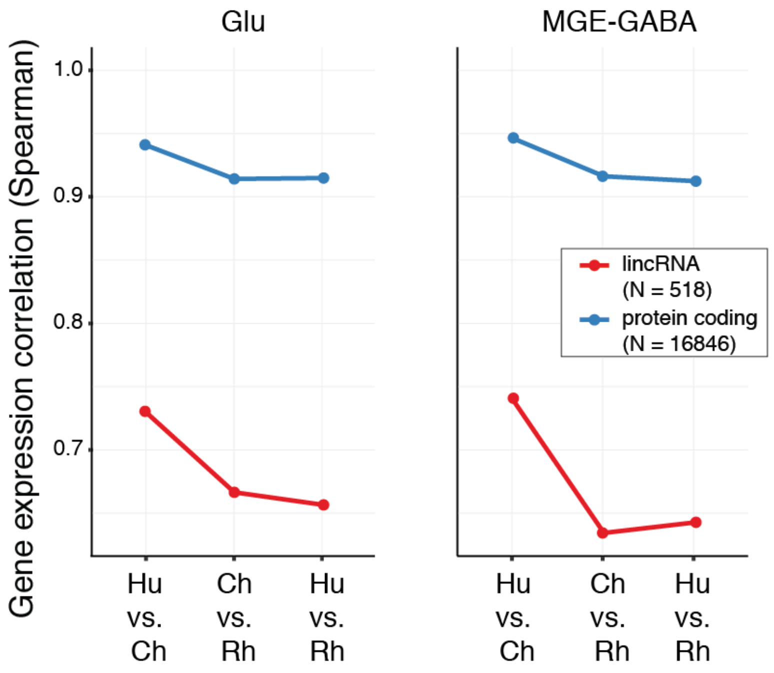

**S3G.** Schematic of the approach to identify species-specific downregulated DA (down-DA) GREs and species-specific downregulated DE (down-DE) genes.

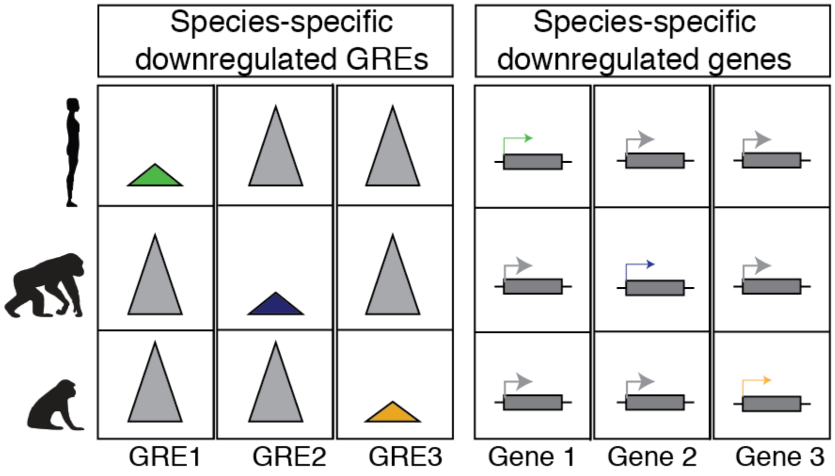


**S3H.** Schematic of the approach to identify concordant pairs of species-specific down-DA GREs and down-DE genes.

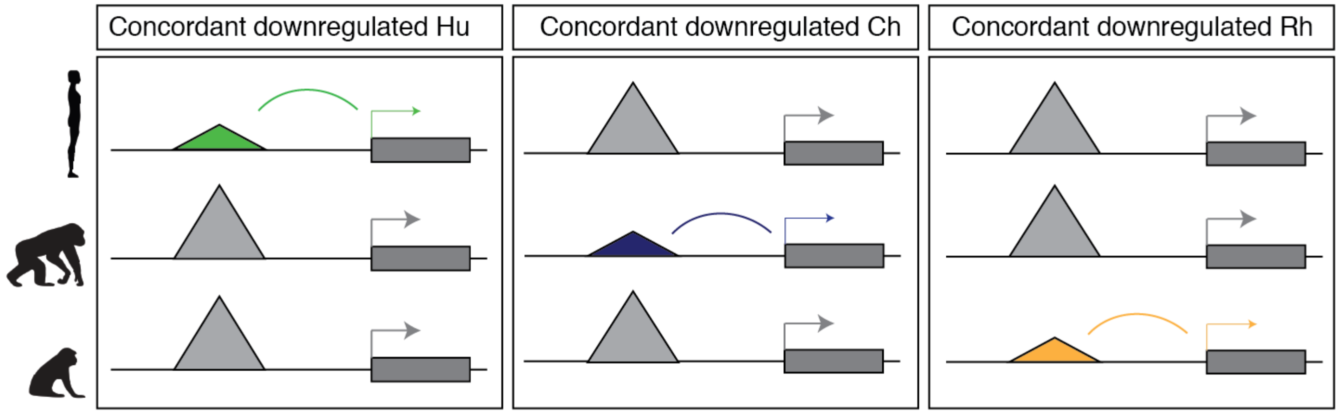

**S3I.** Assessment of the statistical significance of concordant associations between GREs and genes using random shuffling (**see Methods**). For each analysis (Hu-, Ch- or Rh-specific in Glu or MGE-GABA neurons, promoters or enhancers), the assignment of the GRE-gene pairs was randomly shuffled 1000 times. *Gray histograms* show the distributions of concordant GRE counts detected after random shuffling; the *red* dash lines denote the experimentally observed counts. Statistical significance was assessed by t-test. For all analyses, the experimentally observed number of concordant GREs is significantly higher than expected by chance. Shown are the results for species-specific upregulated GREs. Similar results were obtained for downregulated GREs (data not shown).

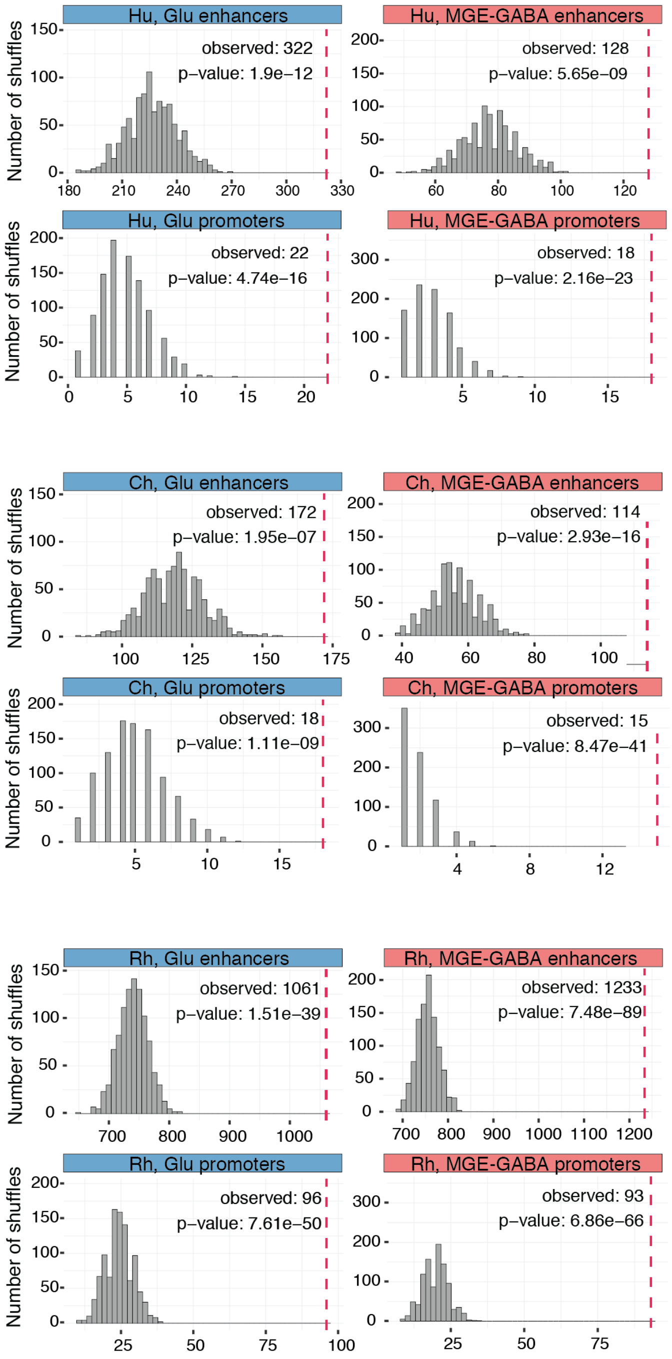

**S3J.** Venn diagram showing the overlaps between the gene sets of Glu- or MGE-GABA-neuronal human-specific up-DE genes which are associated with concordant human-specific up-DA promoters (Prom) or up-DA enhancers (Enh). The majority of human-specific up-DE genes with a concordant up-DA promoter also contain an up-DA enhancer.

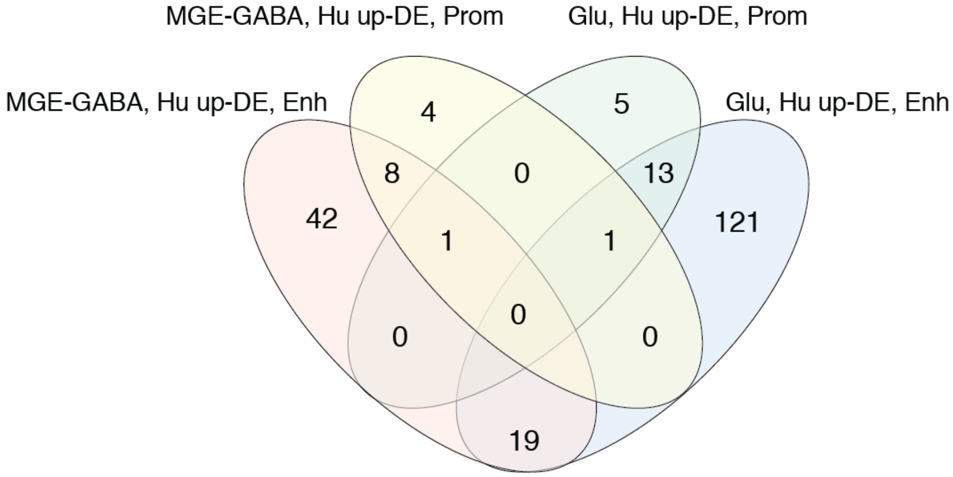


**S3K.** Species-specific up- or down-DE genes in concordant gene-enhancer pairs contain multiple DA enhancers. Shown are fractions of concordant DE genes that are associated with multiple DA enhancers (left panel) and ratios between the numbers of detected DA enhancers and associated concordant DE genes (right panel).

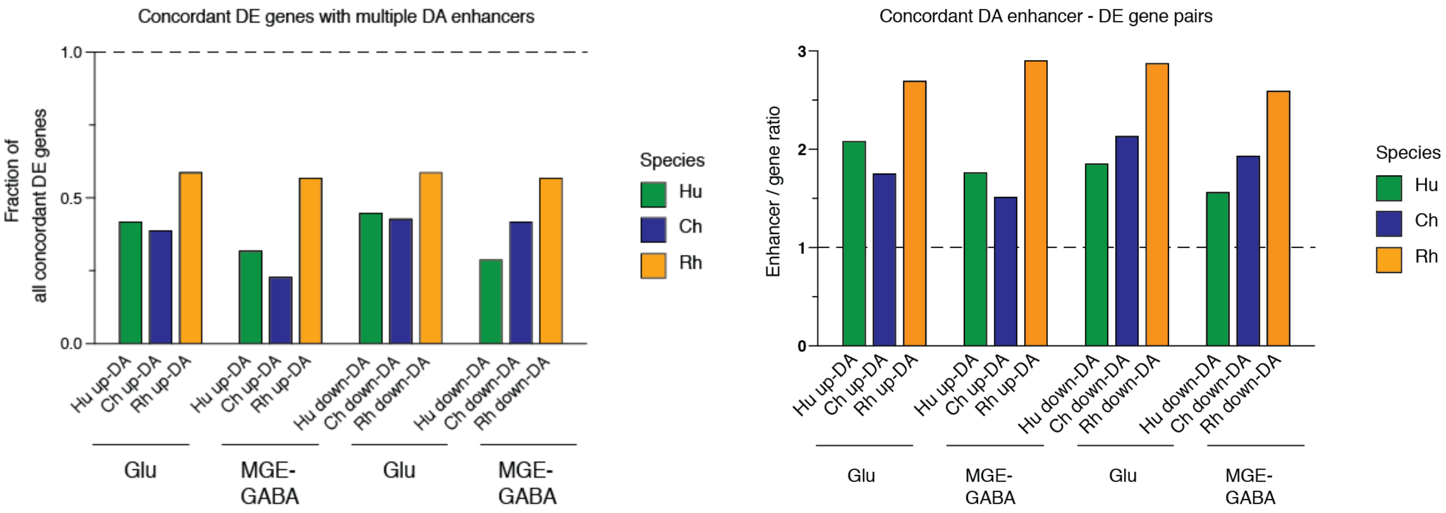

**S3L.** Evolutionary gene expression changes for concordant GRE-gene pairs. Shown are Log2 FC of gene expression between humans and chimpanzees or rhesus macaques for genes associated with DA promoters or enhancers. Evolutionary changes in promoters are associated with larger changes in gene expression compared with evolutionary changes in enhancers. *Left column*: comparison between Hu and Ch, *right column*: comparison between Hu and Rh. *Top row*: data for Glu neurons, *bottom row*: data for MGE-GABA neurons. Statistical significance was assessed by t-test.

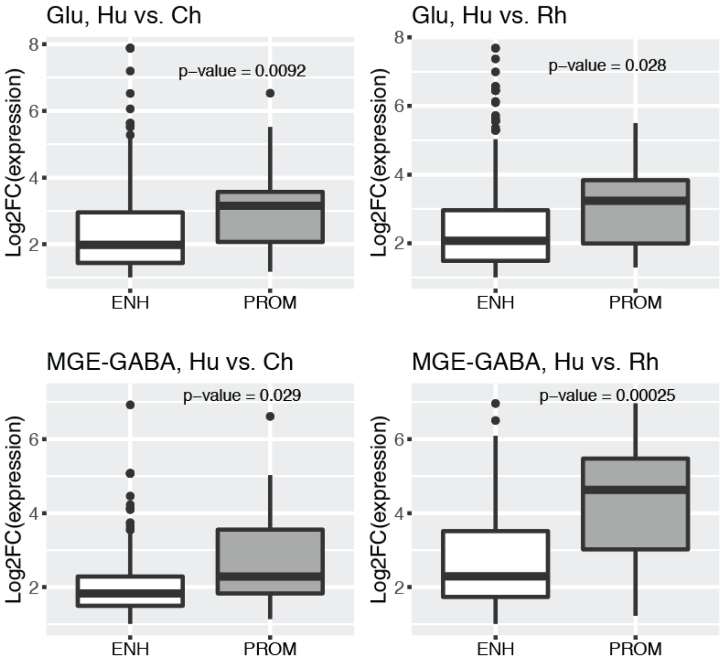

**S3M.** The number of promoters and enhancers within a gene regulatory domain defined by GREAT (**see Methods**) correlates with the gene expression level. Similar results were recently reported in Berthelot et al., 2018. Shown are gene expression levels (average across the three species) as a function of the number of GREs (promoters or enhancers, median across species). *Left column*: data for Glu neurons; *right column*: data for MGE-GABA neurons. *Top row*: data for promoters; *bottom row*: data for enhancers.

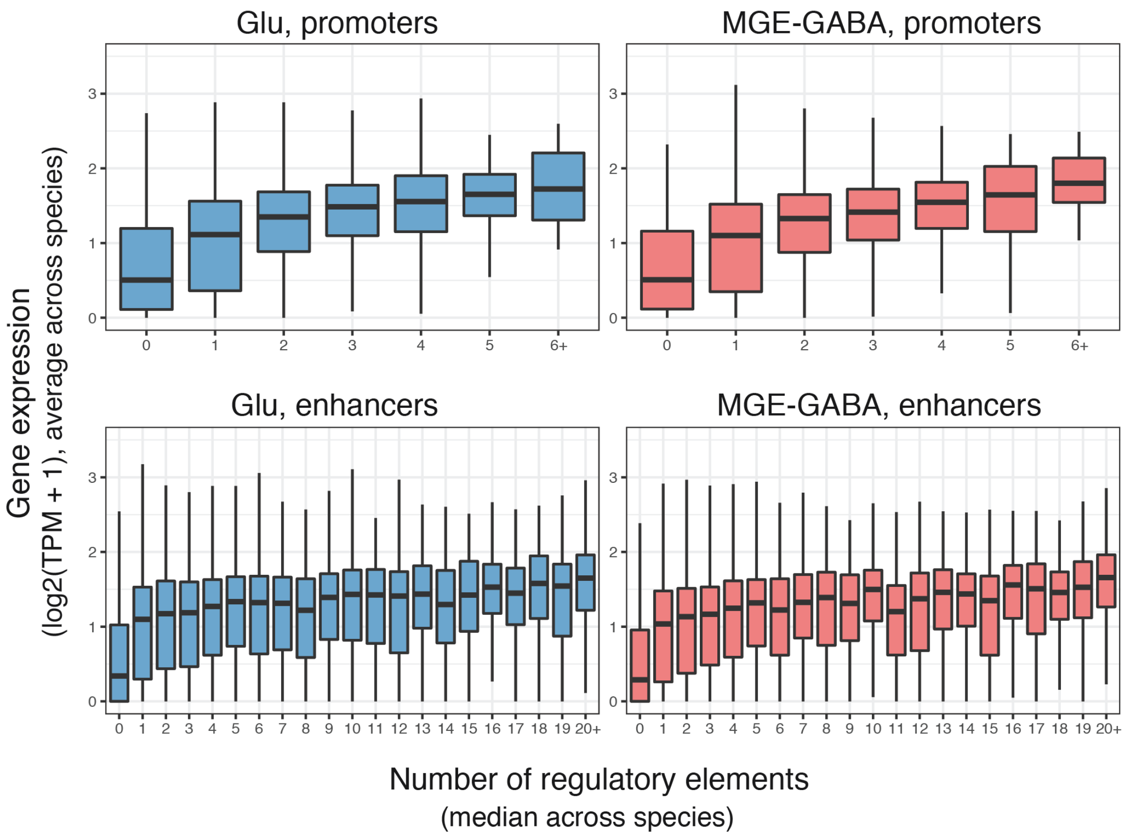

**S3N.** Aggregate quantitative measures of H3K27ac signal intensity for an entire gene regulatory domain correlate with the gene expression level. Two quantitative measures were considered: the sum of H3K27ac signal intensities for all GREs (promoters or enhancers) across a gene regulatory domain (sum intensity, *top row*); the H3K27ac signal intensity for the largest GRE (promoter or enhancer) within a gene regulatory domain (max intensity, *bottom row*). Results are shown separately for promoters and enhancers in Glu or MGE-GABA neurons. Spearman correlation coefficients are shown. All correlations were significant (p-values < 2.2e-16).

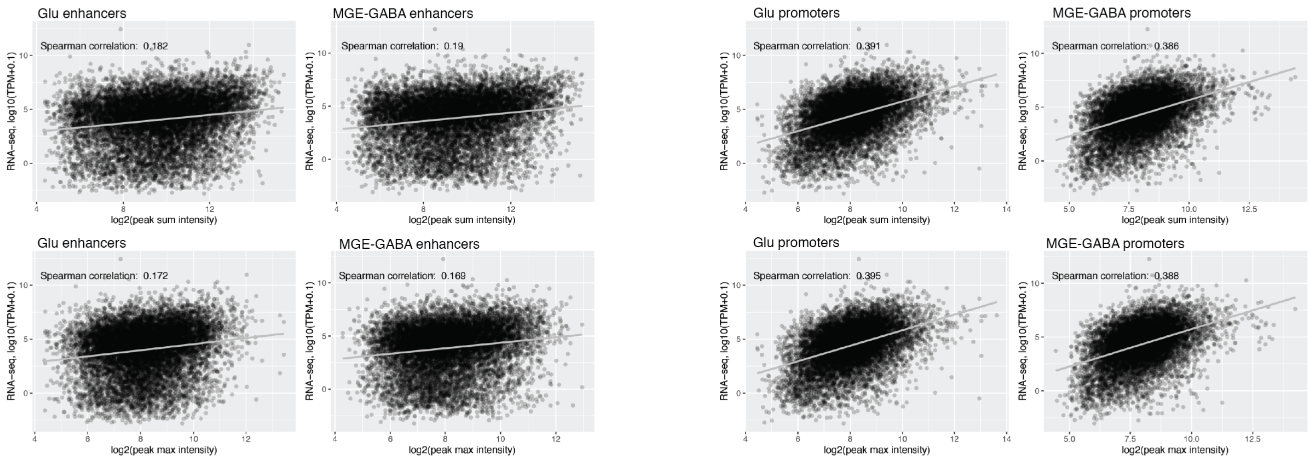


**S3O.** Violin plots showing the relationship between the evolutionary change in the number of DA GREs and the evolutionary change in the expression of the associated genes. Shown is log2 fold change (log2FC) of gene expression (Hu vs. Rh) as a function of the total number of Hu DA GREs minus total number of Rh DA GREs. Data are shown separately for promoters and enhancers in Glu and MGE-GABA neurons. Only genes with significant differences in expression (FDR<0.05) were analyzed.

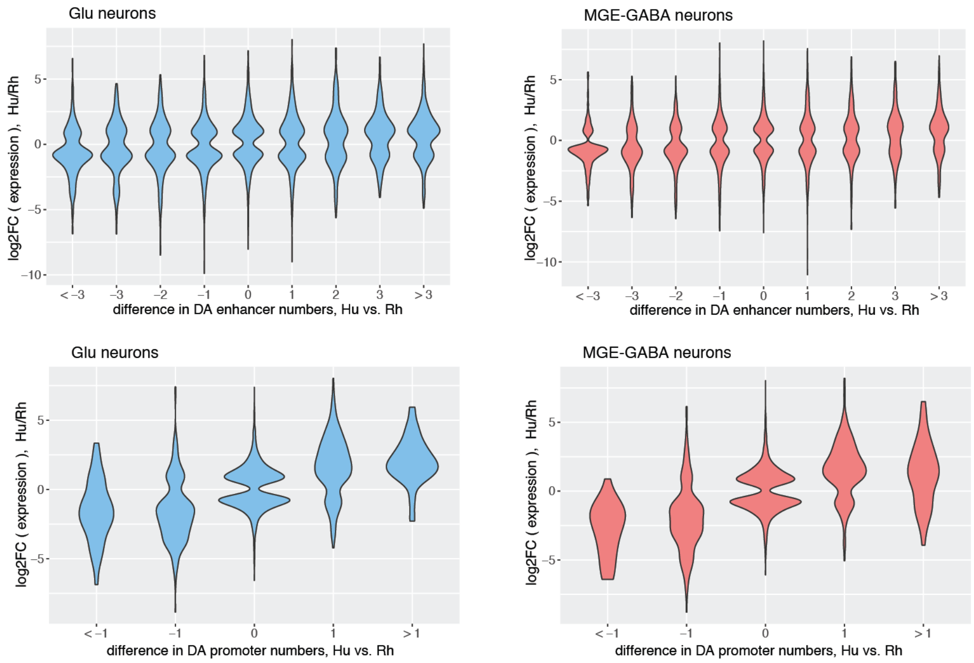

**S3P.** Evolutionary change in aggregate quantitative measures of H3K27ac signal intensity for an entire gene regulatory domain correlates with the evolutionary change in gene expression level. Sum intensity (top row) and max intensity (bottom row) were considered (see **Fig. S3N** for details). All correlations were significant (p-values < 2.2e-16).

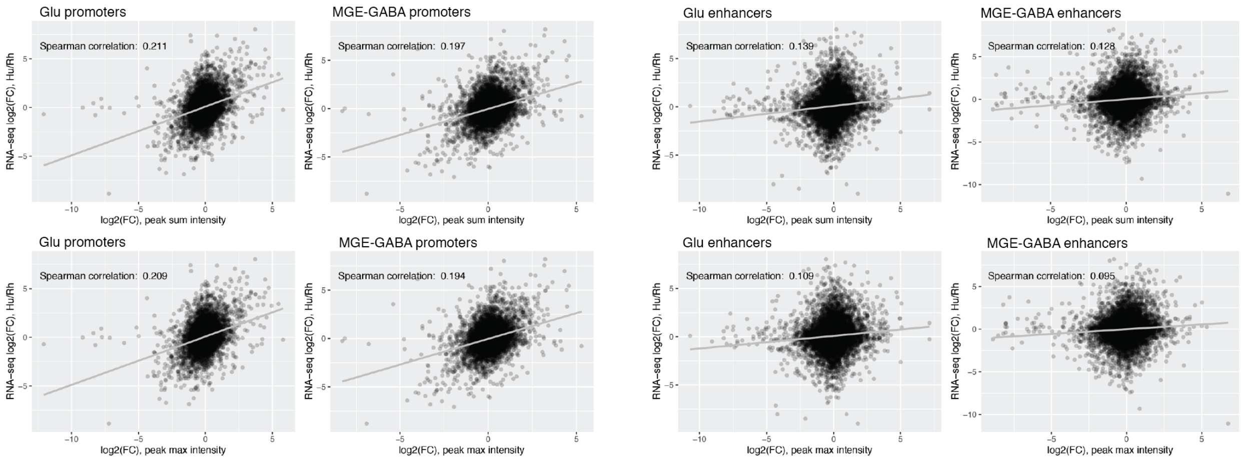

**Figure S4 Human-specific upregulated GREs located near genes implicated in language and neuropsychiatric disorders.**

**S4A.** Human-specific upregulated enhancer within the *FOXP2* gene. Tracks show the H3K27ac signal (RPM-normalized ChIP-seq reads, axis limit 3 RPM) for each cell type (*blue* for Glu neurons and *red* for MGE-GABA neurons) and species. The human-specific enhancer is also Glu-specific (shown in a *dashed box*).

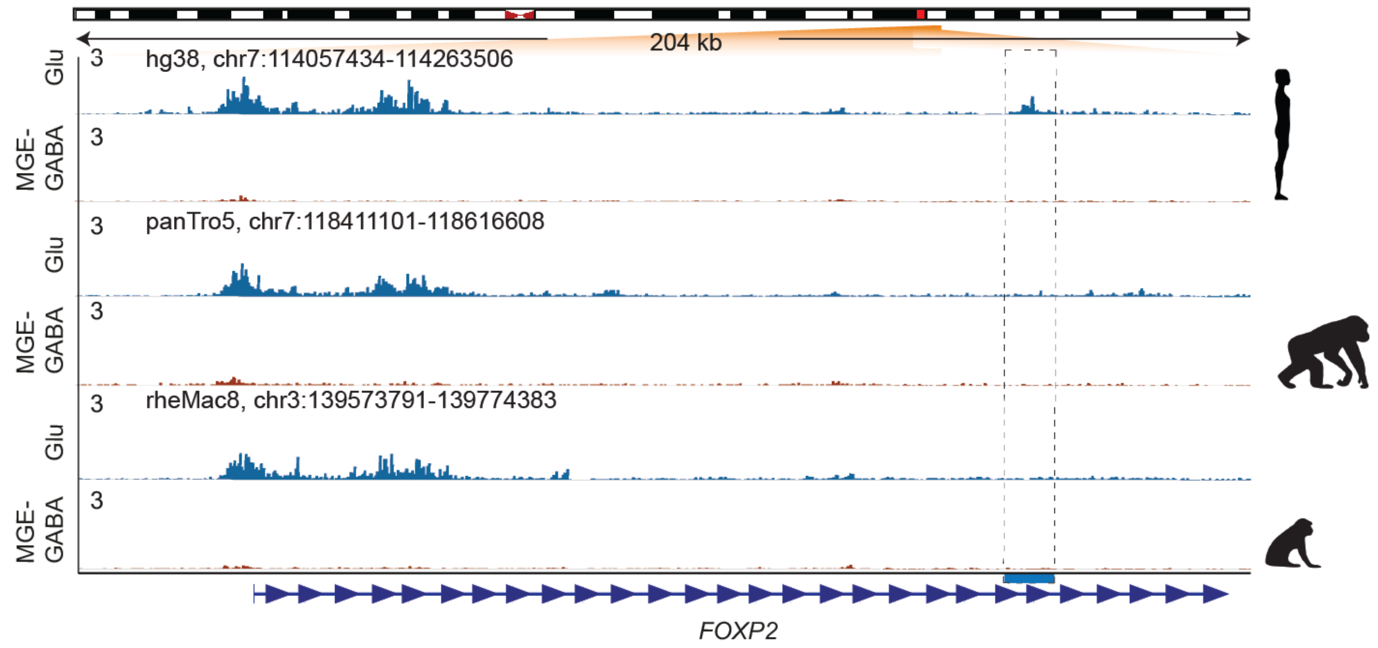


**S4B.** Evolutionary regulatory changes at an enhancer within the *CNTNAP2* locus. Shown are RPM-normalized ChIP-seq reads for each cell type (*blue* for Glu neurons and *red* for MGE-GABA neurons) and species. The depicted human enhancer (*dashed box*) is located within the second intron of *CNTNAP2* and shows human-specific upregulation of the H3K27ac signal in MGE-GABA neurons. ASD-associated SNP rs7794745 (*red arrow*) is harbored within the same enhancer.

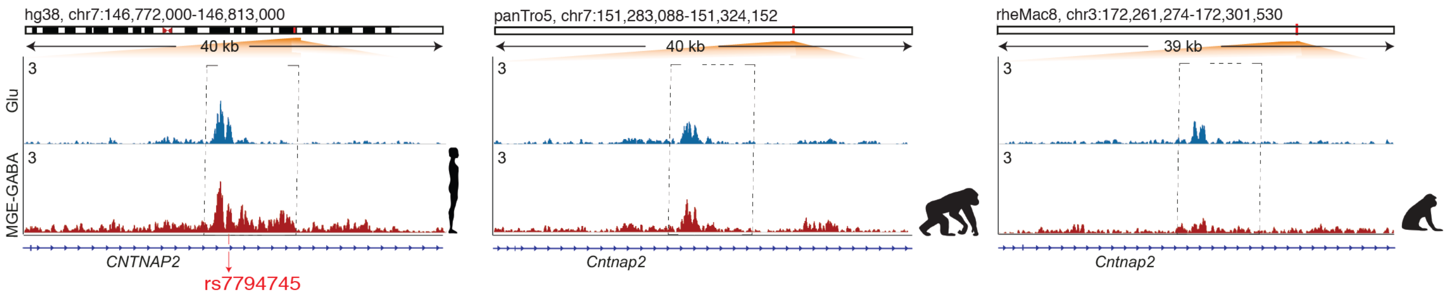

**S4C.** Sequence alignment of primate genomes in the vicinity of human ASD-linked SNP rs7794745. This SNP is located within human-specific upregulated enhancer in MGA-GABA (also see **Fig. 4A**). The major allele nucleotide of *rs7794745* (A>T) is a human-specific substitution compared to other primate species (highlighted).

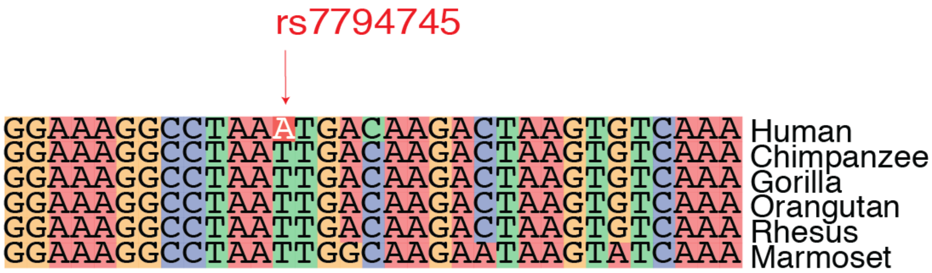

**S4D.** Clusters of human-specific upregulated GREs on chromosome 16.

*Upper panel*: a schematic view of the long (q) arm of chromosome 16 illustrates a non-uniform distribution of human-specific upregulated GREs. The traces depict the locations of annotated genes (*upper trace*), human-specific upregulated GREs detected in Glu neurons (*middle blue trace*) and MGE-GABA neurons (*middle red trace*), as well as all human GREs detected in Glu neurons (*bottom blue trace*) and MGE-GABA neurons (*bottom red trace*). Two Glu-neuronal clusters of human-specific upregulated (up-DA) GREs are shown as *boxed areas*. See **Methods** and **Table 1** for details.

*Bottom panel*: the region of a human chromosome 16 cluster in Glu neurons and the orthologous regions in chimpanzee and rhesus macaque. The cluster is located upstream of the *CDH8* gene. Shown are RPM-normalized ChIP-seq reads (axis limit 3 RPM) for each cell type (*blue* for Glu neurons and *red* for MGE-GABA neurons) and species. Human-specific upregulated enhancers in Glu neurons are highlighted as *dashed boxes*.

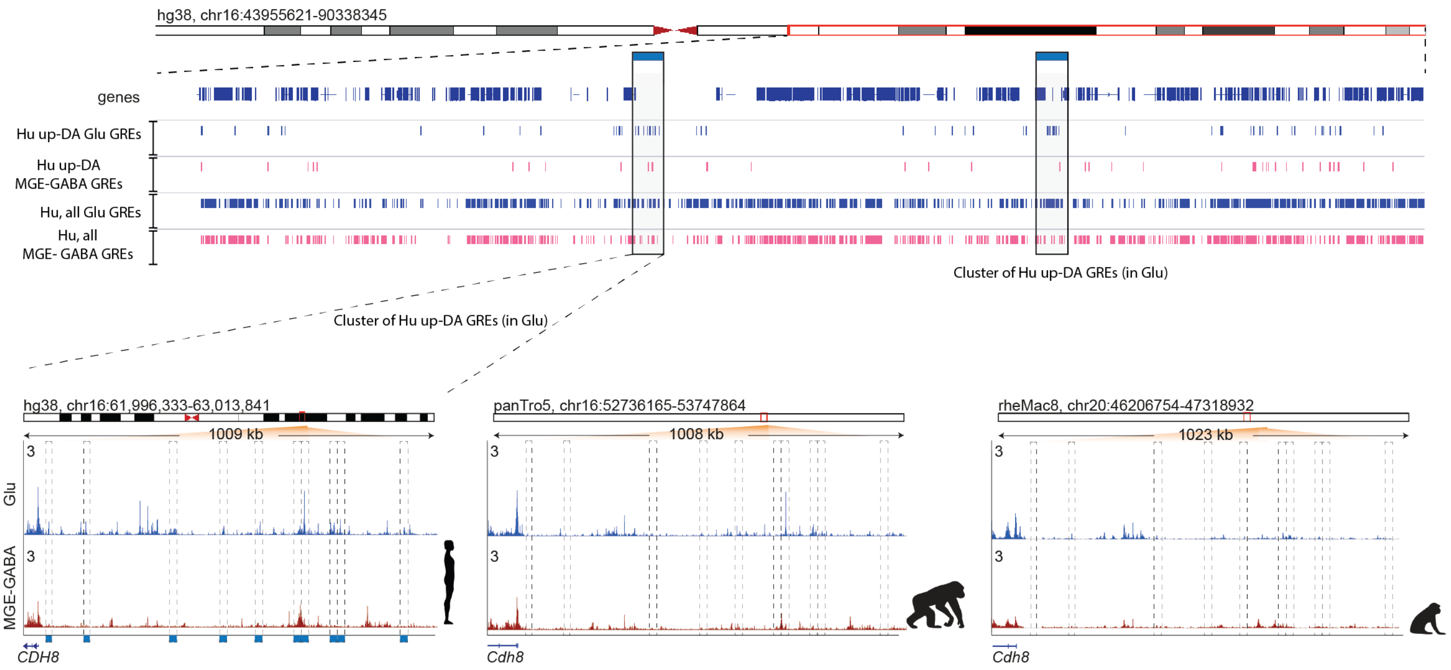

**S4E.** Non-uniform distribution of human-specific upregulated GREs across the genome. The table lists genes or gene sets that lie within genomic regions encompassing clusters of human-specific up-DA GREs. Statistical significance of non-uniform distribution of GREs (see Methods) is shown for each cluster. Clusters detected in Glu and MGE-GABA neurons are highlighted in blue and red, respectively, and the numbers of human-enriched enhancers within each cluster are shown. Also shown are genes linked to ASD according to the SFARI database (56) (*red text*).

| Cell type | chr | Start | End | Minimum FDR per bin | Number of human up-DA peaks | Genes (ASD-associated genes) |
|---|---|---|---|---|---|---|
| Glu | chr2 | 33,200,000 | 35,100,000 | <0.000001 | 18 | *LTBP1; RASGRP3; FAM98A; MYADML* |
| Glu | chr2 | 41,100,000 | 42,200,000 | 0.003701 | 12 | *AC104654.1; C2orf91; PKDCC; EML4* |
| Glu | chr2 | 187,300,000 | 188,500,000 | <0.000001 | 13 | *CALCRL; TFPI; GULP1* |
| Glu | chr2 | 235,800,000 | 236,900,000 | 0.000043 | 12 | *AGAP1; GBX2; ASB18; IQCA1; ACKR3* |
| Glu | chr3 | 99,100,000 | 100,900,000 | <0.000001 | 19 | *COL8A1; CMSS1; TMEM30C; TBC1D23; NIT2; TOMM70A; LNP1; TMEM45A; GRP128; TFG; ABI3BP* |
| Glu | chr6 | 74,400,000 | 75,400,000 | 0.000285 | 11 | *COL12A1; COX7A2; FILIP1* |
| Glu | chr6 | 144,400,000 | 145,700,000 | 0.003798 | 13 | *UTRN; EPM2A* |
| Glu | chr6 | 153,400,000 | 155,000,000 | 0.000001 | 17 | *OPRM1; IPCEF1; CNKSR3; SCAF8* |
| Glu | chr7 | 144,400,000 | 145,400,000 | 0.000523 | 11 | *NOBOX; TPK1* |
| Glu | chr9 | 117,400,000 | 118,400,000 | 0.000099 | 13 | *ASTN2; RP11-500B12.1; TLR4* |
| Glu | chr10 | 9,100,000 | 10,600,000 | 0.000045 | 16 | *LINC00709, CELF2* |
| Glu | chr10 | 48,300,000 | 49,700,000 | <0.000001 | 14 | *MAPK8; ARHGAP22; WDFY4; LRRC18; VSTM4; C10orf128; DRGX; ERCC6; CHAT; C10orf53* |
| Glu | chr14 | 37,500,000 | 38,800,000 | 0.000043 | 13 | *MIPOL1; FOXA1; SSTR1; LINC00639* |
| Glu | chr16 | 62,000,000 | 63,000,000 | 0.000016 | 11 | *CDH8* |
| Glu | chr16 | 76,200,000 | 77,600,000 | 0.000523 | 12 | *CNTNAP4; MON1B; ADAMTS18* |
| Glu | chr18 | 26,600,000 | 27,600,000 | 0.000002 | 11 | *KCTD1; PCAT18; AQP4; CHST9* |
| Glu | chr18 | 31,800,000 | 33,300,000 | 0.000006 | 14 | *TRAPPC8; RNF125; RNF138; MEP1B; GAREM; WBP11P1; KLHL14; CCDC178* |
| Glu | chr18 | 59,400,000 | 60,600,000 | <0.000001 | 14 | *CCBE1; PMAIP1; MC4R* |
| GABA | chr3 | 96,900,000 | 98,000,000 | 0.00214 | 8 | *EPHA6; ARL6; CRYBG3; MINA; GABRR3* |
| GABA | chr5 | 169,100,000 | 170,700,000 | 0.001533 | 10 | *SLIT3; SPDL1; DOCK2; FOXI1; LINC01187; LCP2; KCNIP1* |
| GABA | chr8 | 134,300,000 | 135,400,000 | 0.028364 | 9 | *ZFAT* |
| GABA | chr9 | 95,200,000 | 96,600,000 | 0.00058 | 8 | *FANCC; PTCH1; LINC00476; ERCC6L2; LINC00092; LOC158434; HSD17B3; SLC35D2; HABP4; CDC14B* |
| GABA | chr18 | 53,100,000 | 54,200,000 | 0.00058 | 9 | *DCC; MBD2* |

**S4F.** Evolutionary regulatory changes at the *PENK* (proenkephalin) locus. Representative normalized H3K27ac ChIP-seq tracks for Glu (blue) and MGE-GABA (red) neurons are shown for each species. The dashed box highlights a human- and MGE-GABA-specific enhancer.

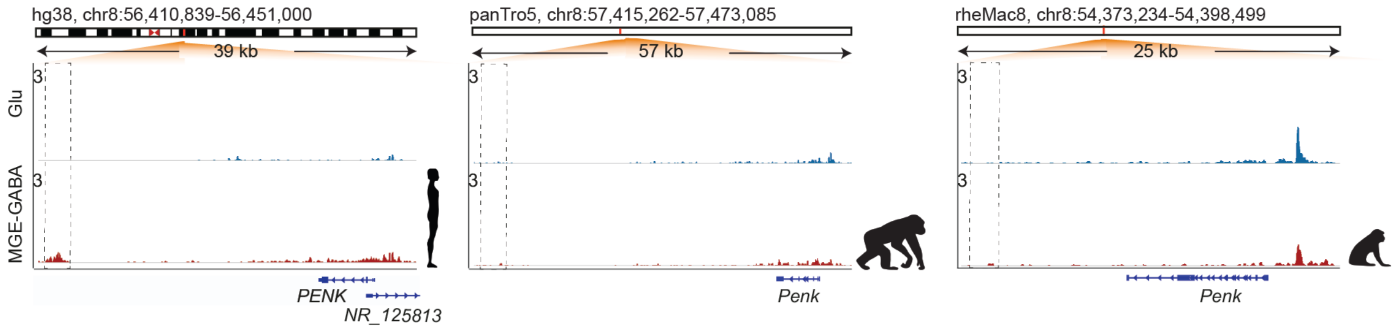


**S4G.** Evolutionary changes in *PENK* gene expression. Shown are TPM-normalized RNA-seq reads for each cell type (*blue* for Glu neurons and *red* for MGE-GABA neurons) and species.

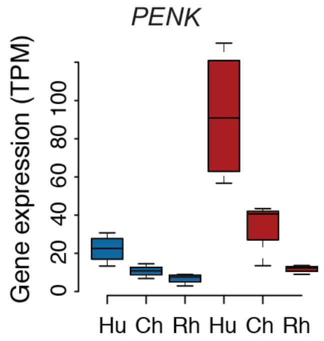
